# Structural and Functional Characterization of G Protein-Coupled Receptors with Deep Mutational Scanning

**DOI:** 10.1101/623108

**Authors:** Eric M. Jones, Nathan B. Lubock, AJ Venkatakrishnan, Jeffrey Wang, Alex M. Tseng, Joseph M. Paggi, Naomi R. Latorraca, Daniel Cancilla, Megan Satyadi, Jessica E. Davis, M. Madan Babu, Ron O. Dror, Sriram Kosuri

## Abstract

In humans, the 813 G protein-coupled receptors (GPCRs) are responsible for transducing diverse chemical stimuli to alter cell state, and are the largest class of drug targets. Their myriad structural conformations and various modes of signaling make it challenging to understand their structure and function. Here we developed a platform to characterize large libraries of GPCR variants in human cell lines with a barcoded transcriptional reporter of G-protein signal transduction. We tested 7,800 of 7,828 possible single amino acid substitutions to the beta-2 adrenergic receptor (β_2_AR) at four concentrations of the agonist isoproterenol. We identified residues specifically important for β_2_AR signaling, mutations in the human population that are potentially loss of function, and residues that modulate basal activity. Using unsupervised learning, we resolve residues critical for signaling, including all major structural motifs and molecular interfaces. We also find a previously uncharacterized structural latch spanning the first two extracellular loops that is highly conserved across Class A GPCRs and is conformationally rigid in both the inactive and active states of the receptor. More broadly, by linking deep mutational scanning with engineered transcriptional reporters, we establish a generalizable method for exploring pharmacogenomics, structure and function across broad classes of drug receptors.

## Introduction

G protein-coupled receptors (GPCRs) are central mediators of mammalian cells’ ability to sense and respond to their environment. The 813 human GPCRs respond to a wide range of chemical stimuli such as hormones, odors, natural products, and drugs by modulating a small set of defined pathways that affect cellular physiology^1^. Their central role in altering relevant cell states makes them ideal targets for therapeutic intervention, with ~34% of all U.S. Food and Drug Administration (FDA)-approved drugs targeting the GPCR superfamily^2^.

Understanding GPCR signal transduction is non-trivial for several reasons. First, GPCRs exist in a complex conformational landscape, making traditional biochemical and biophysical characterization difficult^3,4^. Consequently, most experimentally-determined GPCR structures are truncated, non-native, or artificially stabilized^5^. Even when structures exist, the majority are of inactive states - GPCR conformations that cannot couple with a G protein and cause it to stimulate intracellular signaling. Second, the function of a GPCR depends on its ability to change shape. Static structures from both X-ray crystallography and cryo electron microscopy do not directly probe structural dynamics^6^. Tools such as double electron-electron resonance (DEER) spectroscopy, nuclear magnetic resonance (NMR) spectroscopy, and computational simulation have aided our understanding of GPCR dynamics, but interpreting how structural dynamics relate to function is still difficult^7,8^.

Structure- and dynamics-based analyses generate sets of candidate residues that are potentially critical for function and warrant further characterization. These approaches are complemented by methods that directly perturb protein function such as mutagenesis followed by functional screening. Several reporter gene and protein complementation assays measure GPCR signal transduction by activation of a transcriptional reporter, and are often used to identify and validate important structural residues^9–11^. Such transcriptional reporter assays exist for most major drug receptor classes, including the major GPCR pathways: G_αs_, G_αq_, G_αi/o_, and arrestin signaling^12–14^.

Recent advances in DNA synthesis, genome editing, and next-generation sequencing have enabled deep mutational scanning (DMS) approaches that functionally assay all possible missense mutants of a given protein^15,16^. Several new methods allow for the generation and screening of DMS libraries in human cell lines^17–20^. Function is usually assessed by next-generation sequencing using screens that are bespoke to each gene’s function, or by more general approaches that allow characterization of expression levels rather than function^21^. For GPCRs, the DMS of the CXCR4, CCR5, and T1R2 GPCRs used binding to external epitopes to test expression and ligand binding^22,23^. Unfortunately, such assays tell us little about the signaling capacity of these mutants, which is the primary function of GPCRs and many other drug receptors.

Here we develop an experimental approach to simultaneously profile variant libraries with barcoded transcriptional reporters in human cell lines using RNA-seq. This method is widely applicable to GPCRs and across the druggable genome where transcriptional reporters exist. As a proof-of-principle, we perform DMS on a prototypical GPCR, the β_2_-adrenergic receptor (β_2_AR) and measure the consequences of these mutations through the cyclic AMP (cAMP) dependent pathway.

## Results

### Multiplexed screening platform for G_s_-coupled GPCR signaling

We developed a system to build, stably express, and assay individual variants of the β_2_AR in human cell lines. The β_2_AR primarily signals through the heterotrimeric G_s_ protein, activating adenylyl cyclase upon agonist binding. In our platform, cAMP production stimulates transcription of a barcoded reporter gene, controlled by multimerized cAMP response elements (CRE), which can be quantified by RNA-seq (Fig. 1A). Initially, we generated a HEK293T-derived cell line for stable integration of the GPCR-reporter construct (Fig. 1B, S1A and S1B). We also modified a previously developed Bxb1-landing pad system to allow for stable, once-only integration at the transcriptionally-silent H11 safe-harbor locus to avoid placing the reporter within transcribed genes^24–26^. To prevent endogenous signaling, we knocked out the gene encoding for β_2_AR, *ADRB2*, and verified loss of reporter gene activity in response to the β_2_AR agonist, isoproterenol (Fig. S1C). Our donor vector configuration ensures the receptor and resistance marker are only activated upon successful integration into the landing pad. This vector architecture ameliorates toxicity related to receptor overexpression during library integration (Fig. 1B). Lastly, we included several sequence elements in the donor vector to improve signal-to-noise of the assay: an insulator upstream of the CRE reporter and an N-terminal affinity tag to the receptor (Fig. S1D and S1E). As a result, upon integration of a donor vector expressing wild-type (WT) β_2_AR, isoproterenol induces reporter gene expression in a dose-dependent manner (Fig. 1B).

**Fig. 1.**
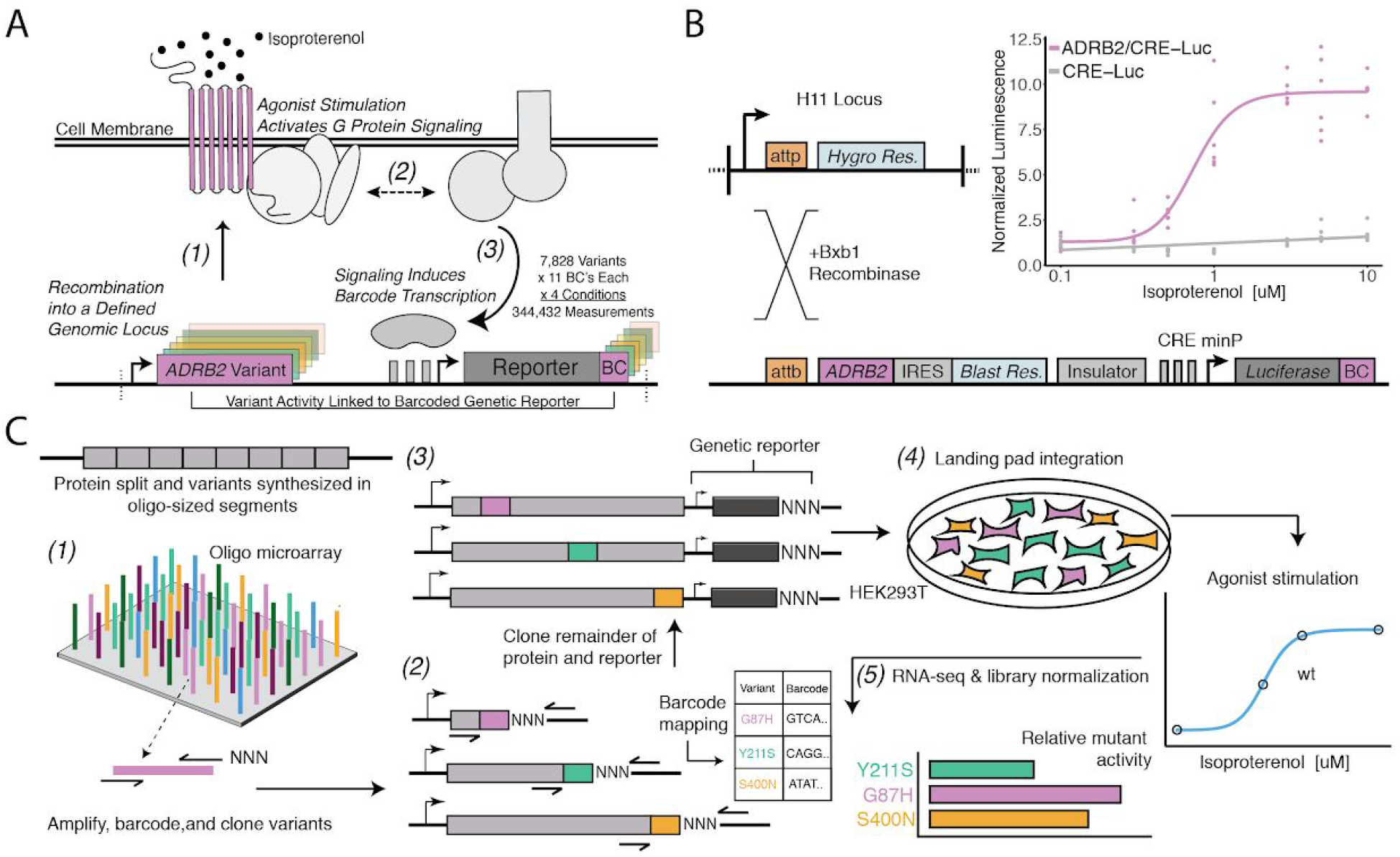
A Platform for Deep Mutational Scanning of GPCRs. **A.** Overview of the Multiplexed GPCR Activity Assay. Plasmids encoding *ADRB2* variants, a transcriptional reporter of signaling activity, and 15 nucleotide barcode sequences that identify the variant are integrated into a defined genomic locus such that one variant is present per cell. Upon stimulation by isoproterenol, G protein signaling induces transcription of the cAMP-responsive genetic reporter and the barcode. Thus, the activity of a given variant is proportional to the amount of barcode mRNA which can be read out in multiplex by RNA-seq. **B.** Schematic detailing the recombination of the reporter-receptor expression plasmid into the landing pad locus. Top Right: Activation of the genetic reporter integrated with (purple) or without (grey) exogenous *ADRB2* into the landing pad when stimulated with isoproterenol in Δ*ADRB2* cells via a luciferase reporter gene assay. **C.** Overview of Library Generation and Functional Assay. Missense variants are synthesized on an oligonucleotide microarray, the oligos are amplified with random DNA barcode sequences appended, and the variants are cloned into wild-type background vectors. Barcode-variant pairs are mapped with next-generation sequencing and the remaining wild-type receptor and genetic reporter sequences are cloned into the vector. Next, the variant library is integrated *en masse* into the serine recombinase (Bxb1) landing pad engineered at the H11 locus of Δ*ADRB2* HEK293T cells. This integration strategy ensures a single pair of receptor variant and barcoded genetic reporter is integrated per cell and avoids crosstalk. After selection, the library is stimulated with various concentrations of the β_2_AR agonist, isoproterenol. Finally, mutant activity is determined by measuring the relative abundance of each variant’s barcoded reporter transcript with RNA-seq.

We designed and synthesized the receptor’s 7,828 possible missense variants in eight segments on oligonucleotide microarrays (Fig. 1C). We amplified the mutant oligos, attaching a random 15 nucleotide barcode sequence, and cloned them into one of eight background vectors encoding the upstream, wild-type portion of the gene. In this configuration, we mapped barcode-variant pairs with next-generation sequencing and subsequently utilized Type IIS restriction enzymes to insert the remaining sequence elements between the receptor and barcode. In the resulting mature donor vector, the barcode is located in the 3’ untranslated region (UTR) of the reporter gene. We integrated the library into our engineered cell line, and developed protocols to ensure proper quantification of library members, most notably vastly increasing the numbers of cells we assayed and RNA processed for the RNA-seq (Fig. S1F and S2A).

### Measurement of mutant activities and comparison to evolutionary metrics

We screened the mutant library at four concentrations of the β_2_AR full-agonist isoproterenol: vehicle control, an empirically determined half-maximal activity (EC_50_), full activity (EC_100_), and beyond saturation of the WT receptor (E_max_). We obtained measurements for 99.6% (7,800/7,828) of possible missense variants (412 residues * 19 amino acids = 7,828 possible missense variants) with two biological replicates at each condition (Fig. 1C). We normalized these measurements against forskolin treatment, which induces cAMP signaling independent of the β_2_AR. Forskolin treatment maximally induces the reporter gene, therefore the relative barcode expression is proportional to the physical composition of the library. Finally, we define activity as the ratio of this value to the mean frameshift (methods). Each variant was represented by 10 barcodes (median), with biological replicates displaying Pearson’s correlations of 0.868 to 0.897 at the barcode level and 0.655 to 0.750 when summarized by individual variants (Fig. S2A).

The heatmap representation of the variant-activity landscape reveals global and regional trends in response to specific subtypes of mutations (Fig. 2A). For example, the transmembrane domain and intracellular helix 8 are more sensitive to substitution than the termini or loops, and this effect becomes more pronounced at higher agonist concentrations (Fig. 2A; all p < 0.001; Mann-Whitney U). The transmembrane domain and intracellular helix 8 are also sensitive to helix-disrupting proline substitutions (Fig. 2B, S2B; all p << 0.001 except TM vs Helix-8; Mann-Whitney U). Microarray-derived DNA often contains single-base deletions that will introduce frameshift mutations into our library^27^. As expected, frameshifts consistently display lower activity than missense mutations regardless of agonist concentration (Fig. 2C; p << 0.001; Mann-Whitney U). Furthermore, the effect of frameshifts are markedly decreased in the C-terminus of the protein (Fig. 2D; p << 0.001; Mann-Whitney U). We also built and integrated previously characterized mutants^28–30^ into our system individually and measured activity with a luciferase reporter gene at the same induction conditions (Fig. 2E and S2C). As expected, known null mutations (D113A and I135W) have significantly diminished activity relative to WT in both systems, even at E_max_ (all p << 0.001; Wald Test). Known hypomorphic mutations (S203A and S204A) also have a significant decrease in activity relative to WT at EC_100_ (all p << 0.001; Wald Test), but are not significantly different than WT at E_max_ as expected (all p > 0.01; Wald Test).

**Fig. 2.**
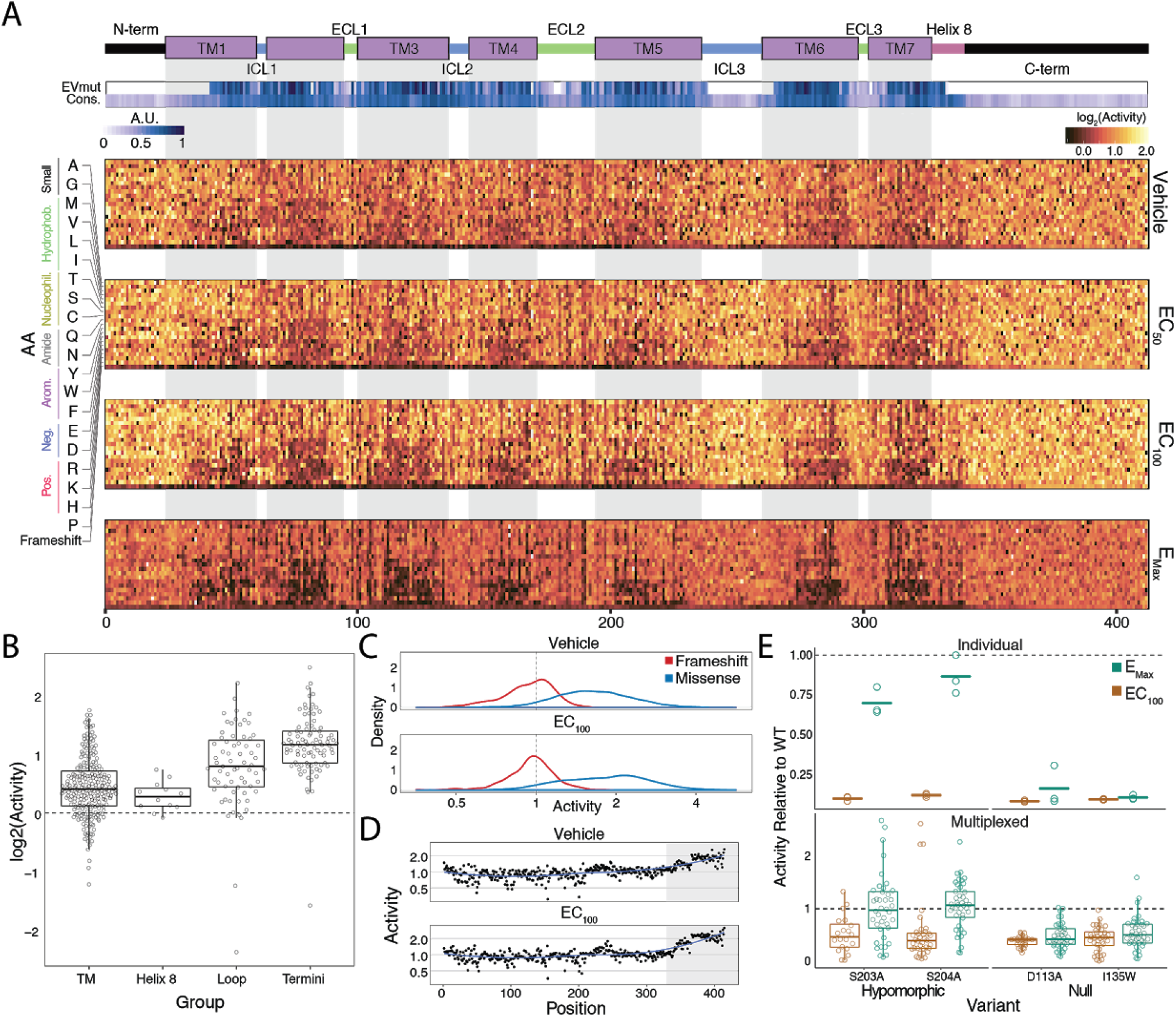
Variant-Activity Landscape for 7,800 Missense Variants of the β2AR and Multiplexed Assay Validation. **A.** Top: Secondary structure diagram of the β_2_AR: the N and C termini are black, the transmembrane helices are purple blocks, and the intra- and extracellular domains are colored blue and green respectively. The EVmutation track (EVmut.) displays the mean effect of mutation at each position as predicted from sequence covariation^33^. Conservation track (Cons.) displays the sequence conservation of each residue across 55 β_2_ AR orthologs^31^. A.U. stands for arbitrary units and the scale for the EVmutation and sequence conservation tracks are individually 0-1 normalized. The shaded guides represent positions in the transmembrane domain. Bottom: The heatmap representation of mutant activity at each agonist condition. Variants are colored by their activity score. relative to the mean frameshift mutation. Activity is the measurement of signaling for each variant relative to the mean frameshift (see methods). **B.** The distribution of mutant activity for proline substitutions is significantly different for amino acids that reside in the transmembrane domain/helix 8 to those in the flexible loops and termini at EC_100_ (all p << 0.001 except TM vs Helix 8; Mann-Whitney U). **C.** The distribution of frameshift mutant activity (red) is significantly different than the distribution of designed missense mutations (blue) across increasing isoproterenol concentrations (both p << 0.001; Mann-Whitney U). Mean frameshift activity marked with a dashed line. **D.** Relative effect of the mean frameshift mutant activity per position is markedly decreased in the unstructured C-terminus of the protein (shaded region), and is consistent across agonist concentration (both p << 0.001; Mann-Whitney U).. Blue line represents the LOESS fit. **E.** Mutant activity measured individually with a luciferase reporter gene compared to the multiplexed assay at EC_100_ and E_Max_ isoproterenol induction. Known null mutations (D113A, I135W) have no dose response between EC_100_ and E_max_ and are significantly different than synonymous mutants at both concentrations in both systems (all p << 0.001; Wald Test). Alternatively, known hypomorphic mutations (S203A, S204A) are significantly different than synonymous mutations at EC_100_ (all p << 0.001; Wald Test), but are not significantly different at E_max_ (all p > 0.01; Wald Test). Bars represent mean value in the luciferase data.

Metrics for sequence conservation and covariation are often used to predict the effects a mutation will have on protein function^31–33^. Mutational tolerance, the mean activity of all amino acid substitutions per residue at each agonist concentration, is highly correlated to conservation, both across species for the β_2_AR (Fig. S3A; Spearman’s ρ = −0.743), and across all Class A GPCRs (Spearman’s ρ = −0.676; Fig. 3A and S3B)^31,33,34^ at EC_100_. Correlation between our data and both predictors increases with agonist concentration up to EC_100_(Fig. S3A and S3B). We found a subset of residues in extracellular loop 2 (ECL2), including C184 and C190 that form an intraloop disulfide bridge, that were more intolerant to mutation than expected given their conservation across Class A GPCRs. This suggests a fairly specific functional role for this motif in the β_2_AR (Fig. 3A). On an individual variant level, mutational responses correlate (Spearman’s ρ = 0.521) with EVmutation, a predictor of mutational effects from sequence covariation (Fig. 3B and S3C)^31,33,34^.

**Fig. 3.**
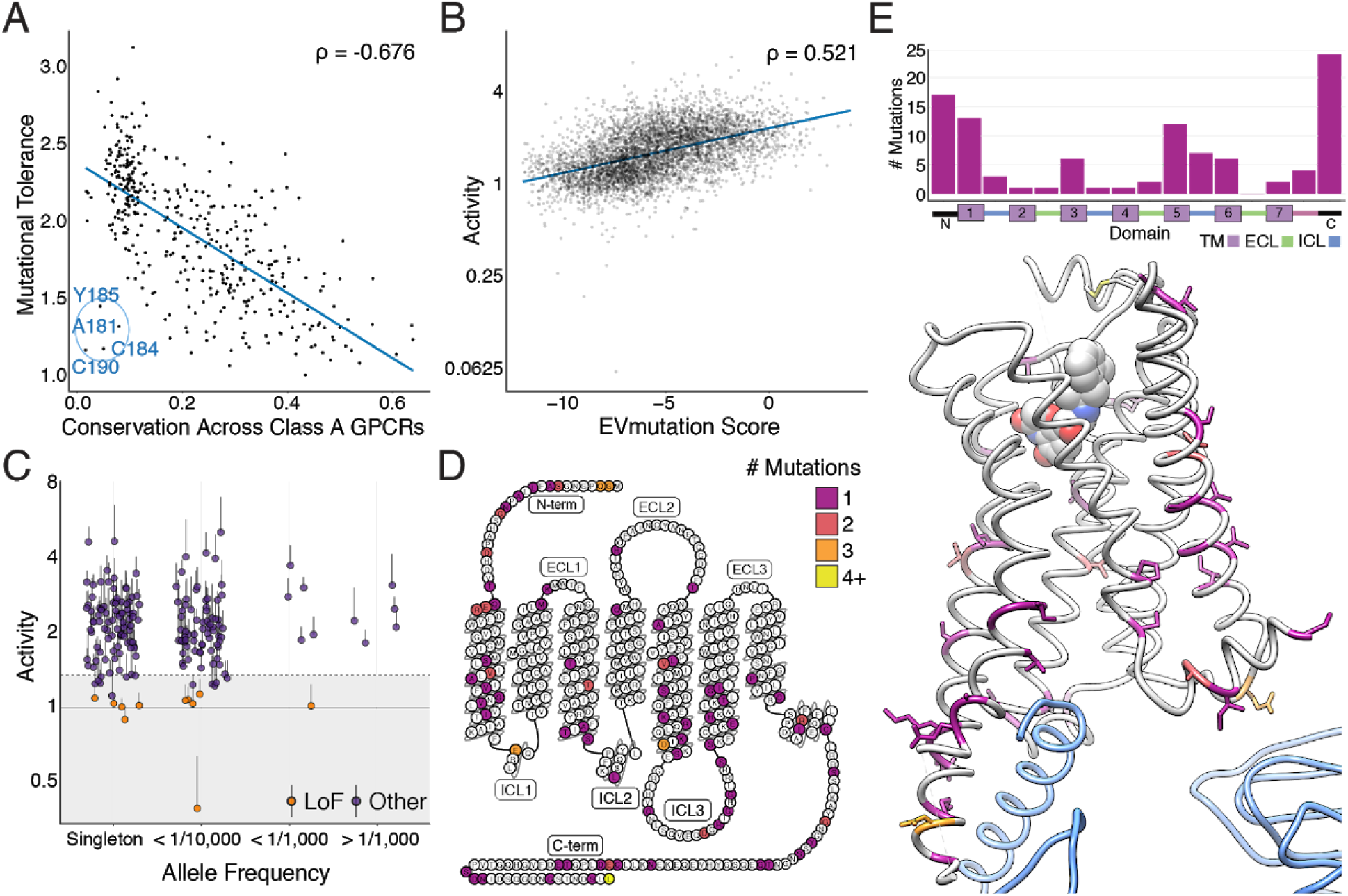
Individual Mutations and Residues Reveal Evolutionary and Structural Insights into β2AR Function. **A.** Positional conservation across Class A GPCRs correlates with mutational tolerance (Spearman’s ρ = −0.676, Pearson’s r = −0.681), the mean activity of all amino acid substitutions per residue at each agonist concentration, at EC_100_. However, four of the least conserved positions (C190, C184, A181, Y185) are highly sensitive to mutation and are located in ECL2, suggesting this region is uniquely important to the β2AR. The blue line is a simple linear regression. **B.** Individual mutant activity correlates with EVmutation (Spearman’s ρ = 0.521, Pearson’s r =0.480) at EC_100_. The blue line is a simple linear regression. **C.** Activity of individual mutants present in the human population from the gnomAD database stratified by allele frequency. Mutations are classified as potential loss of function (LoF) mutations (orange) are classified as such (shaded region) if the mean activity at EC_100_ plus the standard error of the mean (upper error bar) is more than two standard deviations below mean frameshift mutant activity (dashed line). **D.** The distribution of the 100 most basally activating mutations across the β2AR snake plot reveals a clustering of mutants in the termini, TM1, TM5 and TM6. **E.** Top: Distribution of the 100 most basally activating mutations stratified by domain. Bottom: The distribution of the 100 most basally activating mutations across the β2AR 3D structure (PDB: 3SN6). These positions (colored as in D) are concentrated on the surface of the β2AR (G_αs_ shown in blue).

### Population genetics and structural analysis of individual variants

In addition to evolutionary metrics, understanding the functional distribution of *ADRB2* variants found within the human population is important given the extensive variation found among GPCR drug targets^35^. The Genome Aggregation Database (gnomAD) reports variants found across 141,456 individuals^36^, and the majority of the 180 *ADRB2* missense variants are of unknown significance. We classified 11 of these variants as potentially loss of function, by comparing their activity to the distribution of frameshift mutations found in our assay (Fig. 3C; see Methods). Given that measurements of individual mutations are noisy (average coefficient of variation = 0.55), this analysis is best suited as a funnel to guide further characterization (see Discussion).

However, our analysis is more robust when we aggregate the signal of multiple mutations at a given position. Therefore, we compiled a list of the 100 most activating mutations at vehicle control and the 100 least active mutations at EC_100_ and mapped their location on the β_2_AR structure. As expected, the least active mutations tended to reside within the core of the transmembrane domain (Fig. S3D and S3E). Alternatively, the most activating mutations mapped to TM1, TM5, TM6, and residues that typically face away from the internal core of the receptor (Fig. 3D and 3E). Of note, a group of these mutations in TM5 face TM6, which undergoes a large structural rearrangement during receptor activation^37^. Activating mutants are also enriched in the termini, ICL3, and Helix 8. Concentration at the termini is unsurprising, as these regions have known involvement in surface expression^38^. Similarly, the enrichment of activating mutants in ICL3 appears to reflect its role in G protein binding^39–41^. Lastly, we observe a number of activating mutations in the terminal residue, L413. A recent study of genetic variation in human melanocortin 4 (*MC4R*) GPCR also found a gain-of-function mutation at the terminal residue of the receptor, suggesting a possible conserved role for this position in regulating basal activity of GPCRs^42^.

### Unsupervised learning reveals functionally relevant groupings of residues

Given that our data spans thousands of mutations across several treatment conditions, we used unsupervised learning methods to reveal hidden regularities within groups of residues’response to mutation. In particular, we applied Uniform Manifold Approximation and Projection (UMAP)^43^ to learn multiple different lower-dimensional representations of our data and clustered the output with density-based hierarchical clustering (HDBSCAN; Fig. S4)^44^ We found residues consistently separated into 6 clusters that exhibit distinct responses to mutation (Fig. 4A and 4B). Clusters 1 and 2 are globally intolerant to all substitutions, whereas Cluster 3 is vulnerable to proline and charged substitutions. Cluster 4 is particularly inhibited by negatively charged substitutions and Cluster 5 by proline substitutions, while Cluster 6 is unaffected by any mutation. Mapping these clusters onto a 2D snake plot representation shows Clusters 1–5 primarily comprise the transmembrane domain, while Cluster 6 resides in the loops and termini (Fig. 4C). These flexible regions are often truncated before crystal structure determination to minimize conformational variability^45^. Surprisingly, a number of residues from Cluster 5 also map there, suggesting potential structured regions. However, Cluster 5 assignment is largely based on the response of a single proline mutation, and thus is more susceptible to noise than the other clusters (see Discussion).

**Fig. 4.**
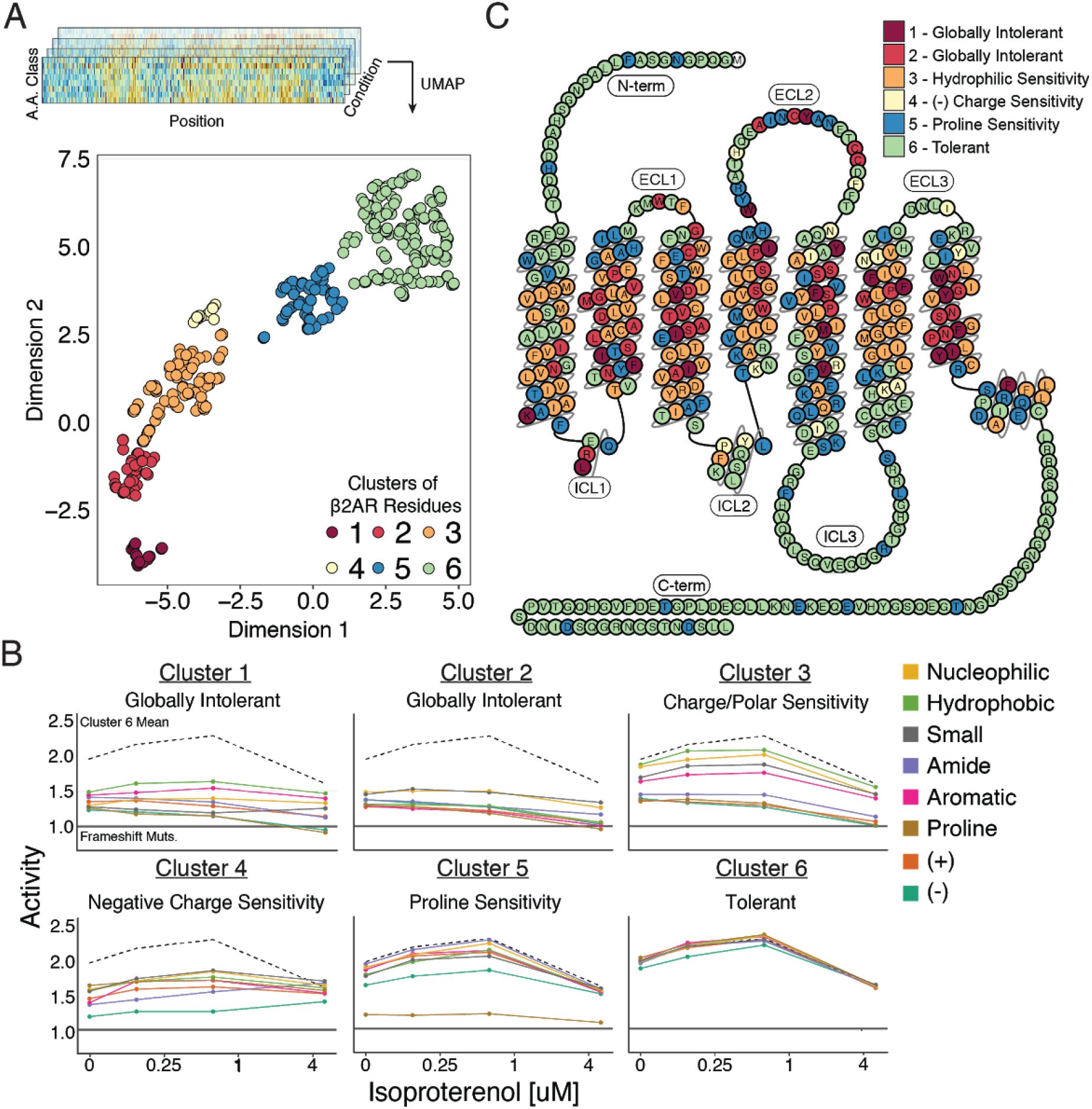
Unsupervised Learning Segregates Residues into Clusters with Distinct Responses to Mutation. **A.** Amino acids were segregated into classes based on their physicochemical properties and mean activity scores were reported by class for each residue. With Uniform Manifold Approximation and Projection (UMAP) a 2D representation of every residue’s response to each mutation class across agonist conditions was learned. Each residue is assigned into one of six clusters using HDBSCAN (see Fig. S5). **B.** Class averages for each of these cluster reveal distinct responses to mutation. The upper dashed line represents the mean activity of Cluster 6 and the lower solid line represents the mean activity of frameshift mutations. **C.** A 2D snake plot representation of β_2_AR secondary structure with each residue colored by cluster identity.

Next, we projected the clusters onto the hydroxybenzyl isoproterenol-bound structure (Fig. S5A; PDB: 4LDL). The globally intolerant Clusters 1 and 2 segregate to the core of the protein, while the charge-sensitive Cluster 3 is enriched in the lipid-facing portion (Fig. S5B). This suggests that differential patterns of response to hydrophobic and charged substitutions could correlate with side chain orientation within the transmembrane domain. Indeed, residues that are uniquely charge sensitive are significantly more lipid-facing than those that are sensitive to both hydrophobic and charged mutations at EC_100_ (Fig. 5A, S5C-D, p = 0.000036; Mann-Whitney U)^46^.

**Fig. 5.**
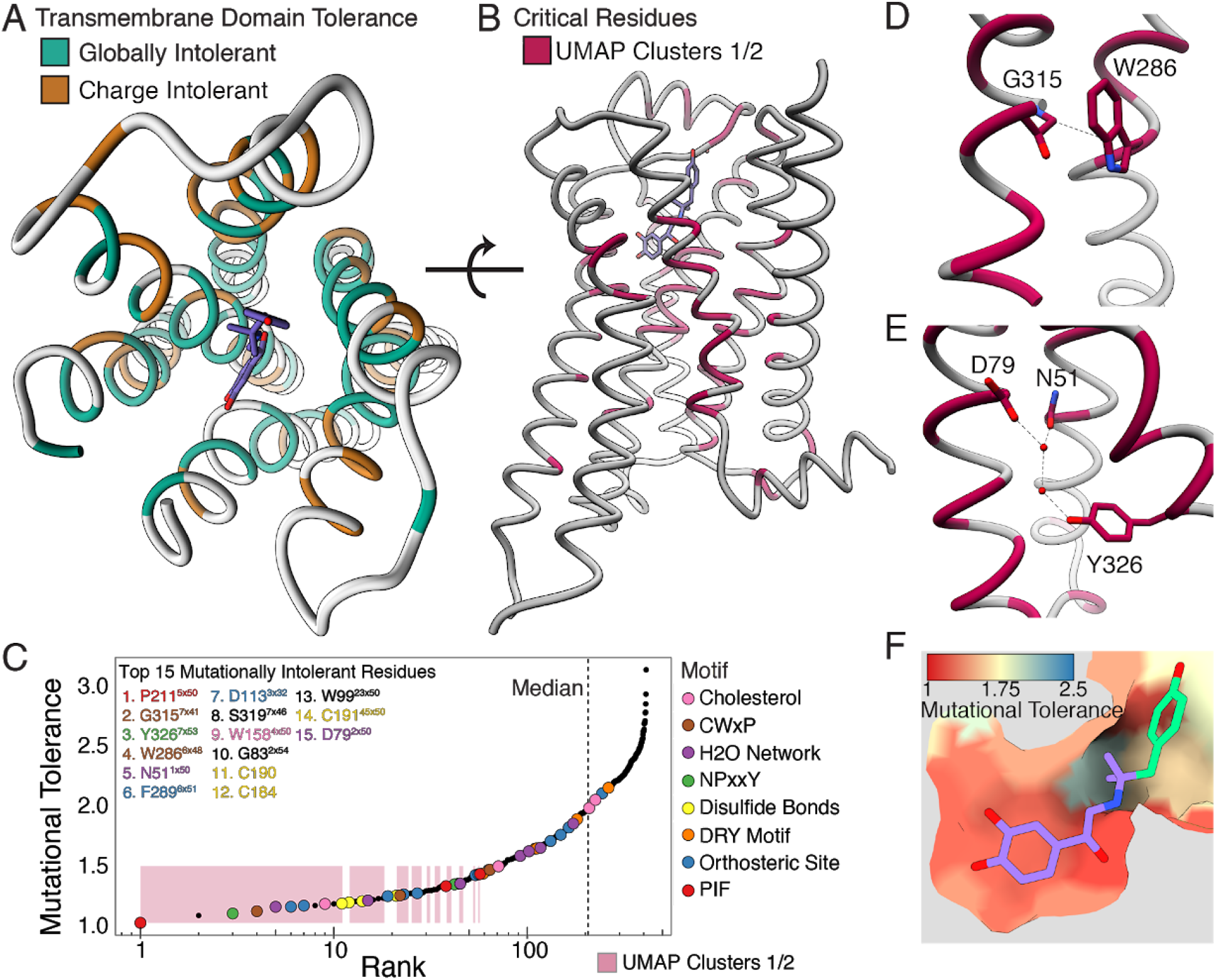
Mutational Tolerance Elucidates Broad Structural Features and Critical Residues of the β_2_AR. **A.** Residues within the transmembrane domain colored by their tolerance to particular classes of amino acid substitution. Teal residues are intolerant to both hydrophobic and charged amino acids (globally intolerant), and brown residues are tolerant to hydrophobic amino acids but intolerant to charged amino acids (charge intolerant). The charge sensitive positions’ side chains are enriched pointing into the membrane, while the globally intolerant positions’ side chains face into the core of the protein (see Fig. S5). **B.** The crystal structure of the hydroxybenzyl isoproterenol-activated state of the β_2_AR (PDB: 4LDL) with residues from the mutationally intolerant Clusters 1 and 2 highlighted in maroon. **C.** 412 β_2_AR residues rank ordered by mutational tolerance at the 0.625 uM isoproterenol condition. Residues in known structural motifs (colored points) are significantly more sensitive to mutation than other positions on the protein (p << 0.001). Dashed line demarcates the median of the ranking. The top 15 mutationally intolerant residues are listed and colored by motif association. **D-F.** Selected vignettes of residues from the mutationally intolerant UMAP clusters and ranking. **D.** W286^6×48^ of the CWxP motif and the neighboring G315^7×41^ are positioned in close proximity. Substitutions at G315^7×41^ are likely to cause a steric clash with W286^6×48^ (PDB: 4LDL). **E.**An inactive-state water-mediated hydrogen bond network (red) associates N51^1×50^ and Y326^7×53^ (PDB: 2RH1). Disruption of this network may destabilize the receptor. **F.** The ligand-bound orthosteric site surface colored by mutational tolerance. Receptor-ligand contacts with the catecholamine head (present in agonist used in assay) are more intolerant to mutation than those in the hydroxybenzyl tail (not present in agonist used in assay) of the isoproterenol analog depicted in this crystal structure (PDB: 4LDL).

### Mutational tolerance stratifies the functional relevance of structural features

Decades of research have revealed how ligand binding is coupled to G-protein activation through a series of conserved motifs^37^. This comprehensive, unbiased screen enables us to systematically evaluate and rank the functional importance of every implicated residue. The globally intolerant UMAP clusters (1 and 2) highlight many residues from these motifs and suggest novel residues for investigation (Fig. 5B). We can further resolve the significance of individual residues within these motifs by ranking the mutational tolerance of positions in these clusters at EC_100_ (Fig. 5C). In fact, 11 of the 15 most mutationally intolerant positions belong to the PIF, CWxP, and NPxxY motif, orthosteric site, a water-mediated bond network, an extracellular disulfide bond, and a cholesterol binding site. Interestingly, the second most intolerant residue is the uncharacterized G315^7×41^ (GPCRdb numbering in superscript^5^). In the active state, G315’s alpha carbon points directly at W286^6×48^ of the CWxP motif, the fourth most intolerant residue, and any substitution at G315^7.x41^ will likely clash with W286^6×48^ (Fig. 5D). We confirmed G315’s intolerance with a luciferase reporter gene assay, where mutants G315T and G315L resulted in complete loss of function (Fig. S6A).

Recent simulations suggest water-mediated hydrogen bond networks play a critical role in GPCR function^47, 7, 48^. The third most intolerant residue in our assay, Y326^7×53^ of the NPxxY motif, is especially important as it switches between two of these networks during receptor activation. In the inactive state, Y326^7×53^ contacts N51^1×50^ and D79^2×50^, two of the top 15 most intolerant positions (Fig. 5E). N51L and N51Y also result in complete loss of function when assayed individually (Fig. S6A). The movement of Y326^7×53^ is also part of a broader rearrangement of residue contacts that are conserved across Class A GPCRs, with the majority of these residues being intolerant to mutation (Fig. S6B)^49^. Aside from G315^7×41^, the other uncharacterized residues in the top 15 include W99^23×50^, S319^7×46^, and G83^2×54^ Given the correlation between mutational tolerance and functional relevance, further investigation of these residues will likely reveal insights into GPCR biology.

Next, we hypothesized residues in the orthosteric site that directly contact isoproterenol would respond uniquely to mutation, however no crystal structure of β_2_AR bound to isoproterenol exists. Using the crystal structure of the β_2_AR bound to the analog, hydroxybenzyl isoproterenol (PDB: 4LDL), we find that residues responsible for binding the derivatized hydroxybenzyl tail have significantly higher mutational tolerance than residues that contact the catecholamine head common to both isoproterenol and hydroxybenzyl isoproterenol at EC_100_ (p = 0.0162; Fig. 5F, Fig. S6C). Given this discrimination, we believe DMS can be a powerful tool for mapping functional ligand-receptor contacts in GPCRs.

GPCR signaling is dependent on a series of intermolecular interactions, and the numerous β_2_AR crystal structures enable us to comprehensively evaluate residues mediating such interactions. For example, cholesterol is an important modulator of GPCR function^50^, and the timolol-bound inactive-state β_2_AR structure elucidated the location of a cholesterol binding site (PDB: 3D4S)^51^. Of residues in this pocket, W158^4×50^ is predicted to be most important for cholesterol binding, and in agreement, W158^4×50^ is the most mutationally intolerant (Fig. S6D). Similarly, a number of studies have mutagenized residues at the G_αs_-β_2_AR interface^11,52–58^, but a complete understanding of the relative contribution of each residue to maintaining the interface is unknown. Most residues are more mutationally tolerant than residues in the intolerant clusters 1 and 2, but the four most intolerant positions are I135^3×54^, V222^5×61^, A271^6×33^, and Q229^5×68^respectively (Fig. S6E). Q229^5×68^ appears to coordinate polar interactions between D381 and R385 of the α5 helix of G_αs_, whereas V222^5×61^ and I135^3×54^ form a hydrophobic pocket on the receptor surface (Fig. S6F).

### A structural latch is conserved across Class A GPCRs

Analysis of the mutational tolerance data has highlighted the functional importance of a previously uncharacterized residues. In particular, W99^23×50^ of extracellular loop 1 (ECL1) is the 13^th^ most intolerant residue, which is unusual as mutationally intolerant residues are rare in the flexible loops. Furthermore, W99^23×50^ is proximal to the disulfide bond C106^3×25^-C191^45×50^, an important motif for stabilization of the receptor’s active state^59,60^. While, aromatic residues are known to facilitate disulfide bond formation, only tryptophan is tolerated at this position^61^. We hypothesize W99’s indole group hydrogen bonds with the backbone carbonyl of the neighboring uncharacterized and mutationally intolerant G102^3×21^, positioning W99^23×50^ towards the disulfide bond. Other aromatic residues are unable to form this hydrogen bond and are less likely to be positioned properly. G102^3×21^ also hydrogen bonds with the backbone amide of C106^3×25^, further stabilizing this region. To verify this claim, we individually confirmed the mutational intolerance of both W99^23×50^ and G102^3×21^ (Fig. S7A).

Interestingly, W99^23×50^, G102^3×21^, and C106^3×25^ are almost universally conserved across Class A GPCRs^62^ (Fig. 6A and S7B). Comparison of over 25 high-resolution structures of class A GPCRs from five functionally different sub-families and six different species revealed that these residues consistently contact each other (Fig. 6B and 6C). Based on the evolutionary and structural conservation across Class A GPCRs, we find W99^23×50^, G102^3×21^, and the C106^3×25^-C191^45×50^ disulfide bond represent a conserved WxxGxxxC motif, forming an extracellular “structural latch” that is maintained consistently throughout GPCRs spanning diverse molecular functions and phylogenetic origins. While a minority of Class A GPCRs lack the Trp/Gly combination of residues in the ECL1 region, these receptors have varying structures in ECL1: an alpha helix (sphingosine S1P receptor), beta strand (adenosine receptor), or even intrinsically disordered (viral chemokine receptor US28) (Fig. S7C).

**Fig. 6.**
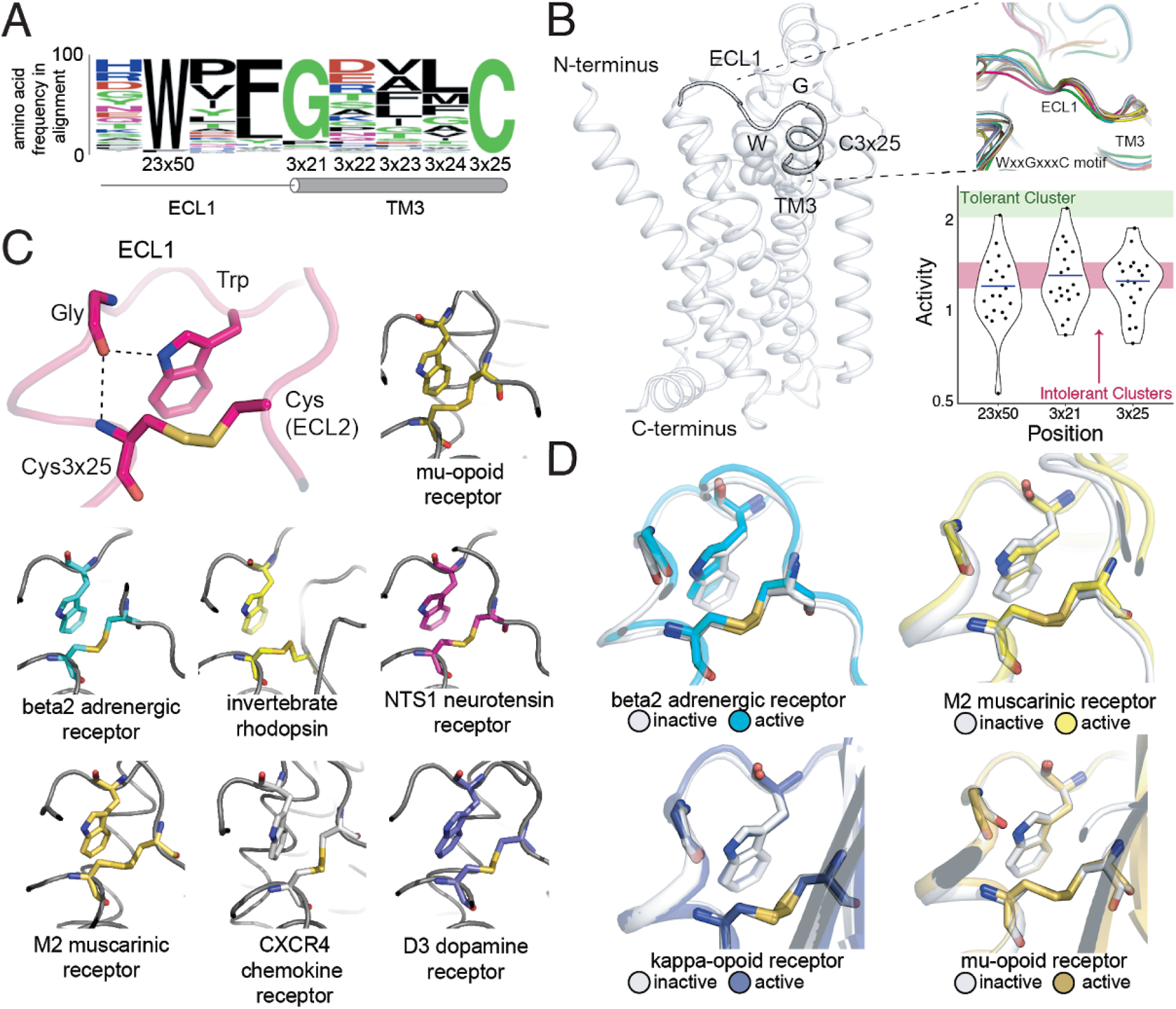
A Conserved Extracellular Tryptophan-Disulfide “Structural Latch” in Class A GPCRs is Mutationally Intolerant and Conformation-Independent. **A.** Sequence conservation of extracellular loop 1 (ECL1) and the extracellular interface of TM3 (202 GPCRs). **B.** Left: Depiction of the interaction of W99^23×50^, G102^3×21^, and C106^3×25^ in ECL1 of the β_2_AR. Top Right: Conservation of the structure of the ECL1 region across functionally different class A GPCRs. Bottom Right: Activity of all 19 missense variants assayed for each of the three conserved residues, with the mean activity (mutational tolerance) shown as a blue bar. The shaded bars represent the mean mutational tolerance ± 1 SD of residues in the tolerant cluster 6 (green) and the intolerant clusters 1 and 2 (red). **C.** A hydrogen bond network between mutationally intolerant positions W99^23×50^, G102^3×21^, and C106^3×25^. Representative examples of the structural latch are shown. **D.** This structural latch is maintained in both the inactive and active state structures for the β_2_AR (inactive: 2RH1, active: 3P0G), the M2 muscarinic receptor (inactive: 3UON, active: 4MQS), the kappa-opioid receptor (inactive: 4DJH, active: 6B73), and the mu-opioid receptor (inactive: 4DKL, active: 5C1M)

To better understand the dynamics of the structural latch, we compared the active and inactive state crystal structures of four representative GPCRs. While the overall RMSD between the inactive and active states for the β2AR, M2 muscarinic receptor, μ opioid receptor and κ opioid receptor are 1 Å,1.5 Å, 1.7 Å and 1.5 Å respectively, the conformation of the latch in the active and inactive states is nearly identical in each receptor (Fig. 6D). This suggests that the extracellular structural latch is part of a larger rigid plug present at the interface of the transmembrane and extracellular regions, which could be important for the structural integrity of the receptor and possibly guide ligand entry.

In Class A receptors lacking components of the WxxGxxxC motif, introducing the Trp-Gly interaction could increase the stability of the receptor for structural studies. In fact, in the BLT1 receptor structure, a Gly mutation at 3×21 was found to be thermostabilizing^63^. Other candidate receptors lacking a Gly at 3×21 include the alpha2B receptor and the neuropeptide FF2 receptor, where the R81G and D112G mutations respectively have potential to increase receptor stability. More broadly, these conserved ECL1/TM3 positions could serve as candidate sites for introducing thermostabilizing mutations.

## Discussion

Our findings showcase a new, generalizable approach for DMS of human protein targets with transcriptional reporters. Such reporters enable precise measurements of gene-specific function that can be widely applied across the druggable genome. We show comprehensive mutagenesis can illuminate the structural organization of the protein and the local environment of individual residues. These results suggest DMS can work in concert with other techniques (e.g. X-ray crystallography, Cryo-EM, and molecular dynamics) to augment our understanding of GPCR structure-function relationships. Moreover, we identify key residues for β2AR function including uncharacterized positions that inform about receptor stability and activation. Importantly, these approaches can be undertaken when direct structural information is unavailable but reporters exist, which is true for most GPCRs.

There are still a number of limitations to our current approach that we expect will improve as we develop the method. Importantly, we did not quantify cell-surface expression directly in our high-throughput functional assay, and thus we cannot distinguish between mutations that substantially affect G protein signaling and those that affect cell-surface expression. In particular, mutations that lead to increased signal in our assays could in fact work by reducing GPCR internalization and not by increasing the intrinsic activity of the receptor. However, we express our variant library in a genomic context at a controlled copy number, dampening the effects of expression-related artifacts typically associated with assays that involve transiently transfected receptor. In addition, expression level alterations can affect the dynamics of signaling and thus may be physiologically relevant. For example, the GPCR *MC4R* is haploinsufficient, and rare heterozygous mutations that eliminate or reduce receptor expression are associated with obesity^42,64,65^. Combining our assay with new generalized, multiplexed assays of protein expression levels in human cells can help tease apart mechanistic reasons for differences in signaling^21^. Secondly, the current signal-to-noise ratio of this approach at single-variant resolution restricted our analyses to mutations with extreme effects on receptor function. This made interpreting single mutations challenging. For example, several mutations within the C-terminus exhibited a sensitivity to proline substitution. This was surprising because the C-terminus is thought to be a flexible, disordered region^66,67^. We individually synthesized and tested three of these mutations (E369P, R253P, and T360P), and found that they did not disrupt function (Fig. S6A). Thus, individual variant data should be confirmed by more traditional assays until the signal-to-noise ratio is improved. However, our measurements are robust in aggregate, and pointed to new receptor biology, providing structural and functional insights. Further improvements to the signal to noise will facilitate the exploration of more subtle aspects of individual mutations.

Looking forward, our method is well-poised to investigate many outstanding questions in GPCR and drug receptor biology. First, individual GPCRs signal through multiple pathways, including pathways mediated by various G proteins and arrestins^68,69–71^. By leveraging transcriptional reporters for each of these pathways, we can understand the mechanisms that underpin signal transduction and biased signaling^72^. Second, GPCRs are often targeted by synthetic molecules with either unknown or predicted binding sites, and often have no known structures. We find ligands imprint a mutational signature on their receptor contacts which could potentially reveal the binding site for allosteric ligands. We also found several regions on the external surface of the receptor where activating mutants are clustered. Since perturbations at these sites appear to increase receptor activity, they could potentially be targeted by positive allosteric modulators or allosteric agonists^50^. Third, the identification of mutations that can stabilize specific conformations or increase receptor expression can aid in GPCR structure determination^73,74^. Fourth, the development of stable cell libraries expressing human medicinally-related GPCR variants can be combined with large-scale profiling against small molecule libraries to build very large-scale empirical maps for how small molecules modulate this broad class of receptors^75–77^. Finally, our approach is generalizable to many classes of drug receptors where transcriptional reporters exist or can be developed^78^, enabling the functional profiling, structural characterization, and pharmacogenomic analysis for most major drug target classes.

## Acknowledgements

We thank the Kosuri Lab for helpful discussions, the UCLA Broad Stem Cell Research Center Sequencing and Flow Cytometry Core, and the Technology Center for Genomics and Bioinformatics for providing next-generation sequencing. We thank Deborah Marks and Jung-Eun Shin for advice implementing EVmutation. We thank Robert J. Lefkowitz and Laura Wingler for suggesting control mutations to test and technical guidance. We thank Hiroaki Matsunami, Joshua S. Bloom, Rishi Jajoo and Rocky O. Cheung for expert technical assistance. Funding: National Science Foundation, Brain Initiative (1556207 to S.K), Ruth L. Kirschstein National Research Service Award (GM007185 to N.B.L.), the USPHS National Research Service Award (5T32GM008496 to E.M.J.), the NIH (DP2GM114829 to S.K.), the Medical Research Council (MC_U105185859 to M.M.B. and A.J.V.), and UCLA. Data and materials availability: Processed data and analysis scripts are available on https://github.com/KosuriLab/b2-dms. Raw data is available with accession number XXXXXX. Plasmids and cell lines are available upon request.

## Author Contributions

E.M.J. and S.K. conceptualized the experiments. E.M.J., D.C., M.S., J.E.D. and J.W. performed the experiments. N.B.L., E.M.J., A.J.V., A.M.T., J.M.P., N.R.L., M.M.B., R.O.D., and S.K. analyzed the results. E.M.J., N.B.L., A.J.V., A.M.T., J.M.P., N.R.L., M.M.B., R.O.D., and S.K. wrote and edited the manuscript.

