## Supplementary Information for "Structural and Functional Characterization of G Protein-Coupled Receptors with Deep Mutational Scanning"

Supplementary Figure 1. Cellular Engineering and Reporter Optimization for Multiplexed Assay
Supplementary Figure 2. Global Metrics of the Multiplexed Screen
Supplementary Figure 3. Correlation with Sequence Conservation and Covariation and Analysis of Individual Mutations
Supplementary Figure 4. Cluster Assignment is Robust Across Different UMAP Embeddings
Supplementary Figure 5. Mutational Profile Suggests Side Chain Orientation and Environment
Supplementary Figure 6. Functional Roles of Mutationally Intolerant Residues
Supplementary Figure 7. The WxxGxxxC Motif is Highly Conserved Across Class A GPCRs

Supplementary Table 1. Processed Data
Supplementary Table 2. Mutational Tolerance
Supplementary Table 3. Primers Used in this Study

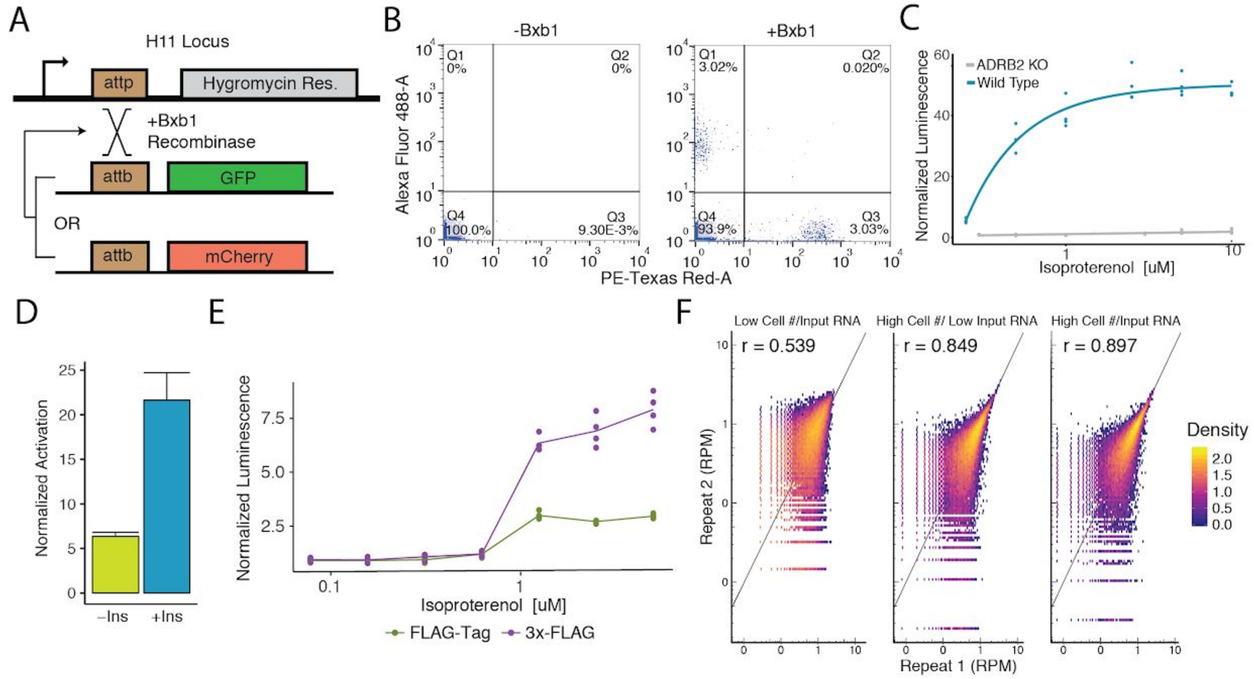


**Supplementary Figure 1. Cellular Engineering and Reporter Optimization for Multiplexed Assay. A.** Schematic of experiment to ensure the landing pad is present at single copy in the genome and thus recombine a single donor plasmid per cell. Single copy integration is essential to prevent receptors of variable functionality to activate barcoded reporters mapped to other variants. Upon co-expression of the promoterless GFP and mCherry plasmids with bxb1 recombinase sites, a cell line with a single landing pad will exclusively integrate one cassette. Therefore, cells will be either GFP$^{+}$ or mCherry$^{+}$ but never both. **B.** Flow Cytometry plots detailing the percentage of GFP$^{+}$ and mCherry$^{+}$ cells when transfected with an equimolar ratio of promoterless GFP and mCherry expression cassettes with or without Bxb1 recombinase expression. **C.** Activation of a cAMP-responsive luciferase reporter gene integrated in the landing pad when stimulated with isoproterenol in a WT or ΔADRB2 background. Activation of the reporter in the WT background emphasizes the importance of generating a Δ*ADRB2* cell line for the multiplexed assay. **D.** Activation of the genetic reporter/*ADRB2* expression cassette with or without a CHS4 DNA insulator upstream of the reporter gene integrated in the landing pad when stimulated with isoproterenol. **E.** Fold activation of an integrated genetic reporter/*ADRB2* expression cassette with a FLAG-tag or 3x-FLAG tag fused to the N-terminus of *ADRB2*. **F.** Alterations to the processes in the multiplexed assay that improve barcode abundance estimate. Initially, we seeded 2,320,000 cells/replicate and processed ~25 ug of total RNA per replicate and observed modest correlation (r = 0.539). We then scaled up to seeding 32,378,680 cells/replicate and correlation improved (r = 0.849). Finally, we began processing ~650 ug of total RNA/replicate and noticed further improvement (r = 0.897).

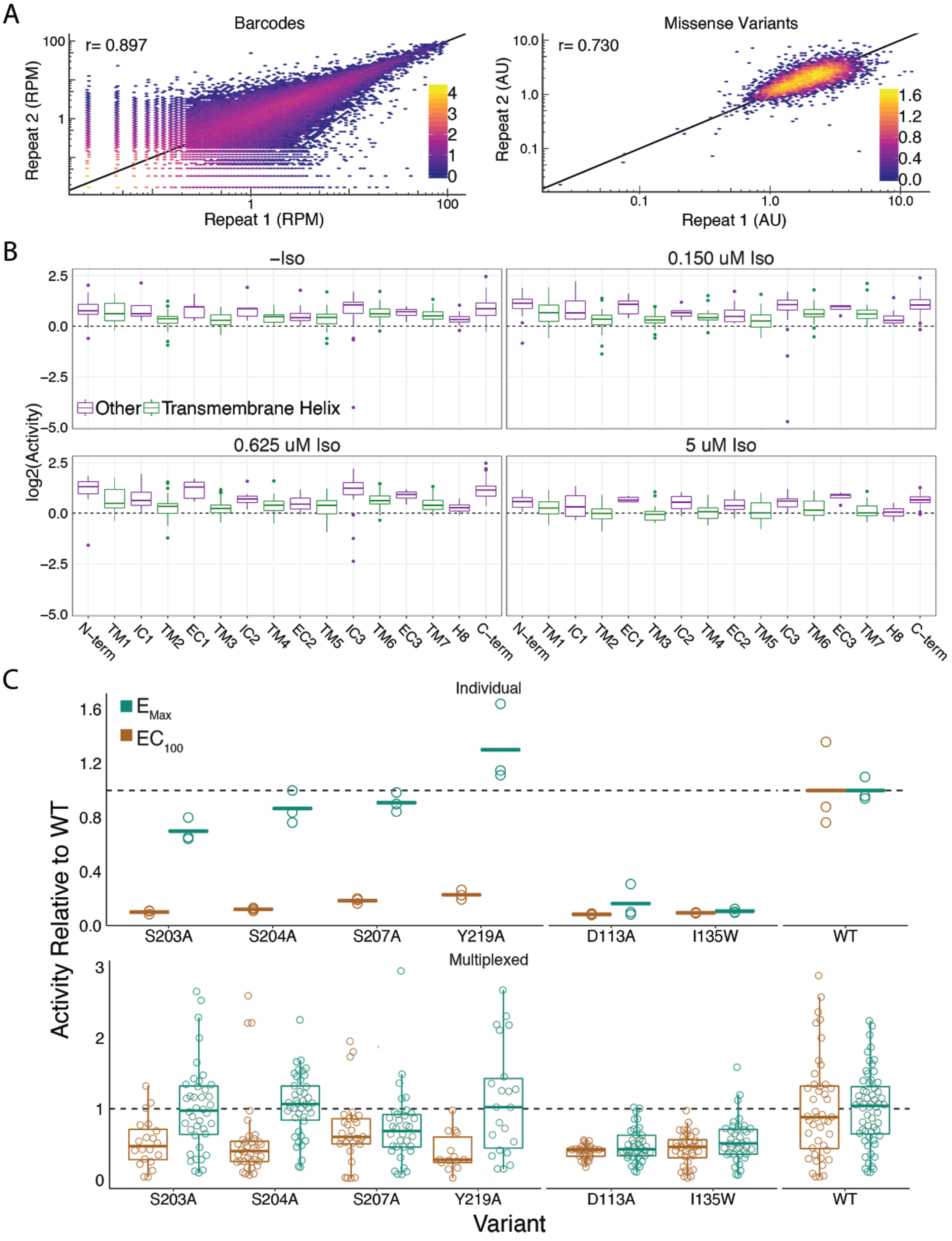
A
Barcodes
r= 0.897
Repeat 2 (RPM)
Repeat 1 (RPM)
Missense Variants
r= 0.730
Repeat 2 (AU)
Repeat 1 (AU)
B
-Iso
0.150 uM Iso
0.625 uM Iso
5 uM Iso
Other
Transmembrane Helix
log2(Activity)
N-term
TM1
IC1
TM2
EC1
TM3
IC2
TM4
EC2
TM5
IC3
TM6
EC3
TM7
H8
C-term
C
Individual
E Max
EC 100
Multiplexed
Activity Relative to WT
S203A
S204A
S207A
Y219A
D113A
I135W
WT
Variant

**Supplementary Figure 2. Global Metrics of the Multiplexed Screen. A.** The measurements between barcodes at the RNA-seq level are correlated ($r = 0.889$, $r = 0.893$, $r = 0.897$, $r = 0.868$) at all agonist concentrations (0, 0.150, 0.625, and 5 uM Iso). Similarly, the mean forskolin-normalized values for each variant are correlated at every concentration as well ($r = 0.655$, $r = 0.687$, $r = 0.730$, $r = 0.750$). Representative plots from 0.625 uM Iso shown. Bars represent log10 counts per hex-bin. **B.** Activity of proline mutations stratified by domain reveals a proline sensitivity in the transmembrane domain across all agonist conditions. **C.** Mutant activity measured individually with a luciferase reporter gene compared to the multiplexed assay at $EC_{100}$ and $E_{Max}$ isoproterenol induction. Known hypomorphic mutations (S203A, S204A, S207A, Y219A) are significantly different than WT at $EC_{100}$ (all $p \ll 0.001$ except S207A in multiplex - $p = 0.009$; Wald Test), but recover to near WT activity at $E_{max}$ (all $p > 0.01$; Wald Test) in both systems. Alternatively, known null mutations (D113A, I135W) are significantly different than WT regardless of the drug concentration (all $p \ll 0.001$; Wald Test). Bars represent mean value in luciferase data.

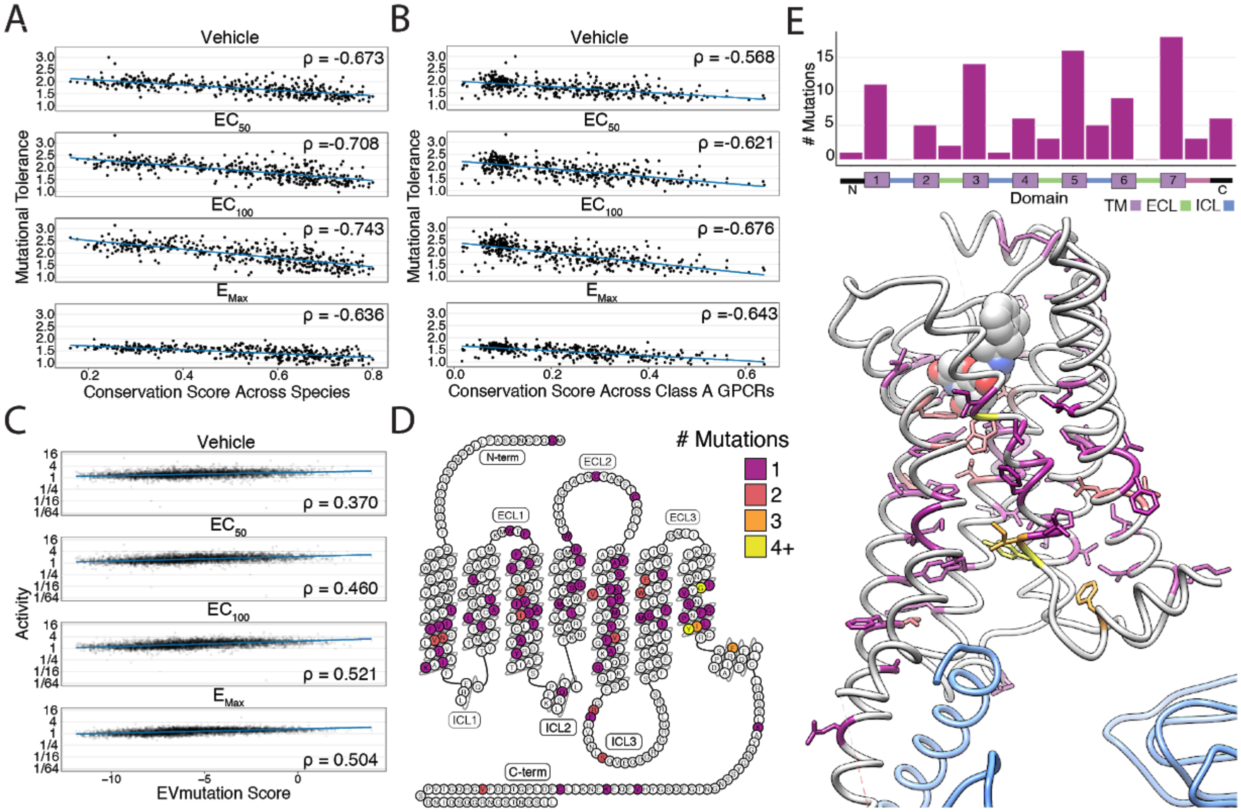


**Supplementary Figure 3. Correlation with Sequence Conservation and Covariation and Analysis of Individual Mutations. A.** Mutational tolerance is highly correlated with species-level sequence conservation and is maximized at $EC_{100}$ (Spearman's ρ = -0.673, Pearsons r = -0.654; ρ =-0.708, r = -0.694; ρ = -0.743, r = -0.729; ρ = -0.636, r = -0.622; for -Iso, 0.150 uM Iso, 0.625 uM Iso, and 5 uM Iso, respectively; all p << 0.0001). Here we calculated sequence conservation using the Jensen-Shannon divergence from a multiple alignment of 55 ADRB2 orthologs from the OMA database. The blue line is a simple linear regression. **B.** Similarly, mutational tolerance of the individual positions is highly correlated with sequence conservation across Class A GPCRs and is also maximized at $EC_{100}$ (ρ = -0.568, r = -0.566; ρ =-0.621, r = -0.631; ρ = -0.676, r = -0.681; ρ = -0.643, r = -0.653 for -Iso, 0.150 uM Iso, 0.625 uM Iso, and 5 uM Iso, respectively; all p << 0.0001). The blue line is a simple linear regression. **C.** Activity measurements for individual substitutions correlates with predictions from EVMutation, and is maximized at $EC_{100}$ (ρ = 0.370, r = 0.327; ρ = 0.460, r= 0.413; ρ = 0.521, r = 0.480; ρ = 0.504, r = 0.488; all p << 0.0001). The blue line is a simple linear regression. **D.** The location of the 100 most deleterious mutations by activity score at $E_{Max}$ (5 uM) Iso on the β2AR snake plot. Mutations are concentrated in the transmembrane domain. **E.** Top: Distribution of the 100 most deleterious mutations by activity score at $E_{Max}$ (5 uM) Iso stratified by domain. Bottom: Location of these mutants on the 3D structure of the β2AR. These positions (colored as in D) tend to face into the core of the $β_2$AR (PDB: 3SN6; Gs in blue).

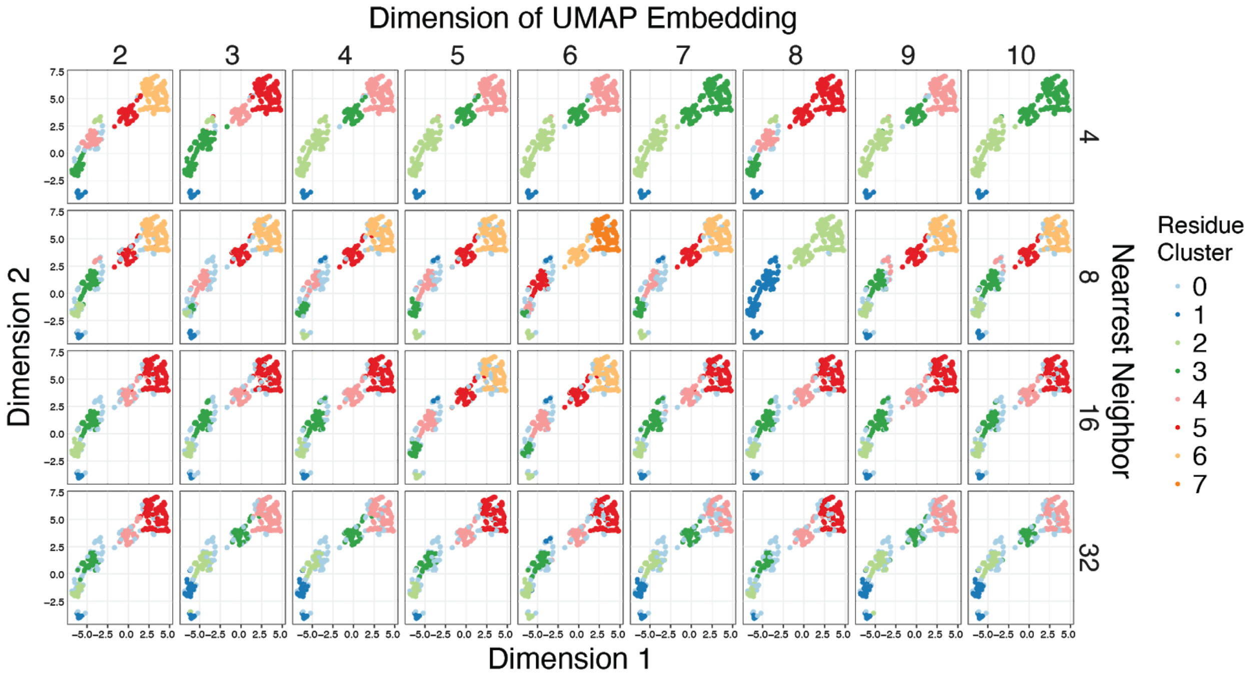


**Supplementary Figure 4. Cluster Assignment is Robust Across Different UMAP Embeddings.** Given the high dimensionality of the mutational responses, Uniform Manifold Approximation and Projection (UMAP)[1] was used to learn lower-dimension representations of the all the mutational data across agonist conditions summarized by amino acid class before clustering the output with HDBSCAN (minimum cluster size = 10)[2]. To ensure that the clustering results are not biased by a particular UMAP embedding, a hyperparameter search was run over the dimension and nearest neighbor parameters of UMAP. The HDBSCAN cluster assignments were plotted on a 2D UMAP embedding to ease visualization. Points that HDBSCAN does not assign to a cluster are colored powder blue. Groups of residues reliably cluster together regardless of the UMAP embedding, and residues were assigned to one of six distinct clusters.

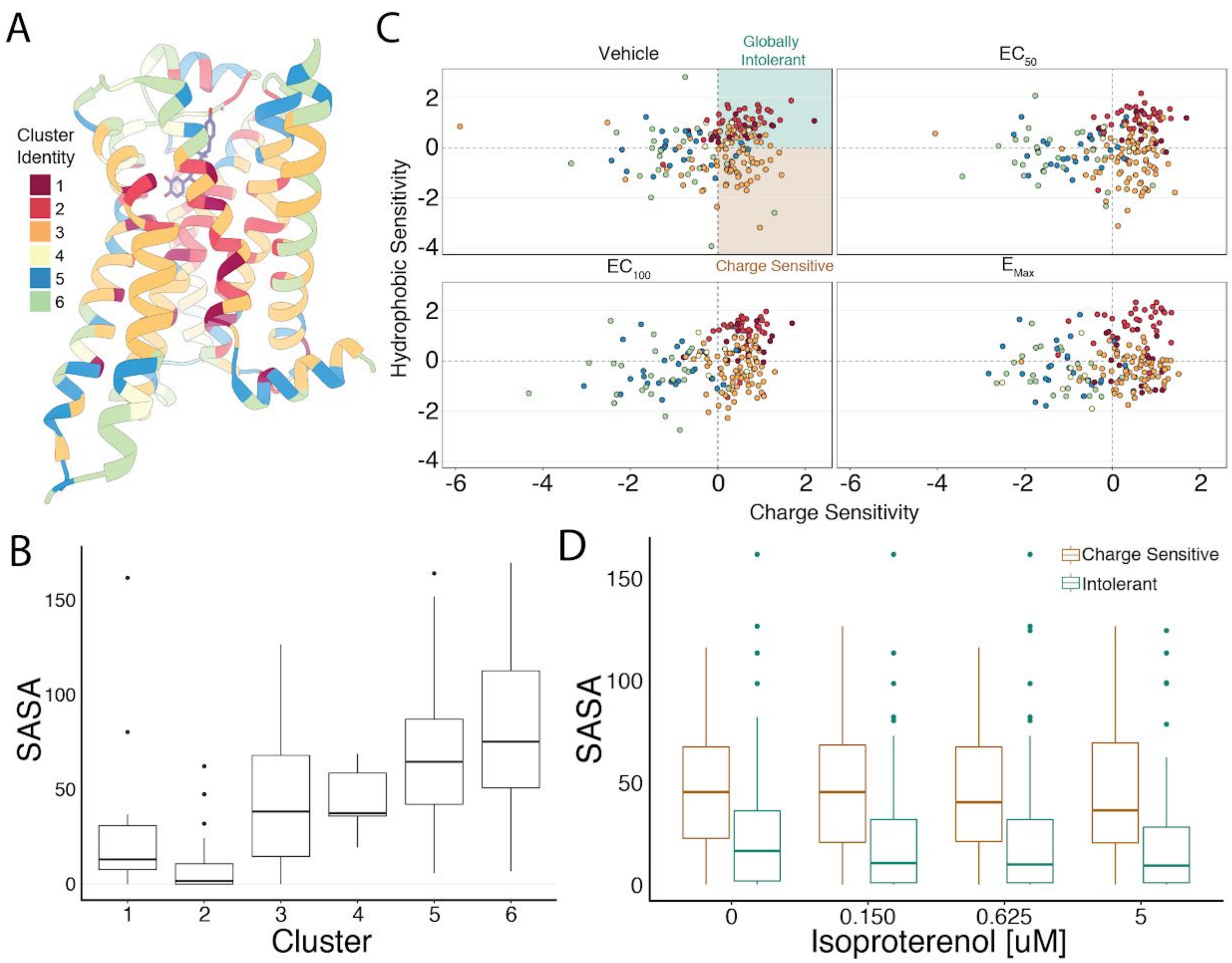


**Supplementary Figure 5. Mutational Profile Suggests Side Chain Orientation and Environment. A.** The crystal structure of the hydroxybenzyl isoproterenol-activated state of the $\beta_2$AR (PDB: 4LDL) with residues colored by UMAP cluster identity. **B.** Distributions of Solvent Accessible Surface Area (SASA) for each cluster at $EC_{100}$. **C.** Hydrophobic versus Charge Sensitivity across all drug conditions. Points are colored by cluster identity. Residues are defined as globally intolerant to substitution if their Hydrophobic and Charge Sensitivity is greater than 0. Similarly, residues are defined to be uniquely charge sensitive if their Hydrophobic Sensitivity is less than 1 and their Charge Sensitivity is greater than 1 (see Methods). **D.** The median SASA is significantly higher for positions in the charge sensitive clusters when compared to the ones in the intolerant clusters across all drug concentrations (all $p < 0.0005$). This suggests that the charge sensitive cluster residues point towards the lipid whereas the ones that are intolerant tend to be buried in the core of the protein.

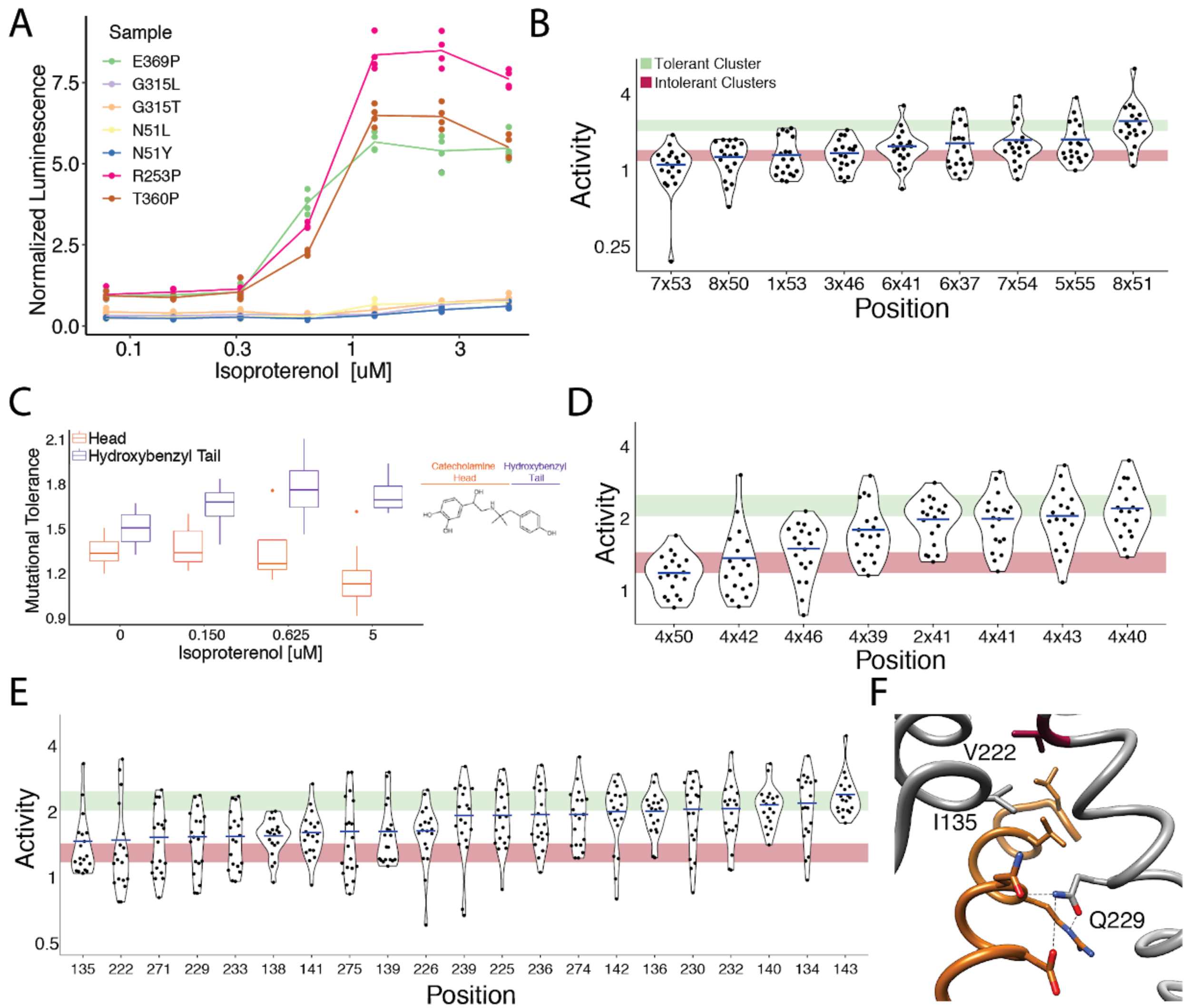


**Supplementary Figure 6. Mutational Intolerance of Functionally Related Residues. A.** Relative activation of an integrated CRE luciferase reporter gene for β2AR missense variants mentioned in the manuscript. **B.** Functional consequences of mutation for a set of residues near the G protein coupling region involved in GPCR activation[3]. **C.** Residues that interact with the catecholamine head (orange) of hydroxybenzyl isoproterenol have significantly lower mutational tolerance than those that interact with the hydroxybenzyl functional group on the tail (purple). In our assay, we screened with isoproterenol lacking the hydroxybenzyl tail. These differences are significantly different at $EC_{50}$ ($p = 0.028$), $EC_{100}$ ($p = 0.016$), and saturating agonist concentration ($p = 0.008$). **D.** Functional consequences of mutation at predicted contacts of a cholesterol binding pocket determined in the timolol-bound structure of the $\beta_2$AR inactive state (PDB: 3D4S). As predicted, the highly conserved W158$^{4x50}$ is the most constrained residue. The shaded bars represent ± 1 standard deviation of the mutational tolerance in the tolerant Cluster 6 (green) or the intolerant clusters 1 and 2 (red). The mean activity of every mutation at a given position (the mutational tolerance) is shown as a blue bar. **E.** Effects of mutations at residues in the Gs interface. **F.** Three of the four most intolerant $\beta_2$AR residues at the G protein interface (brown) from the $\beta_2$AR-Gs complex crystal structure (PDB: 3SN6), V222$^{5x61}$, I135$^{3x54}$, and Q229$^{5x68}$.

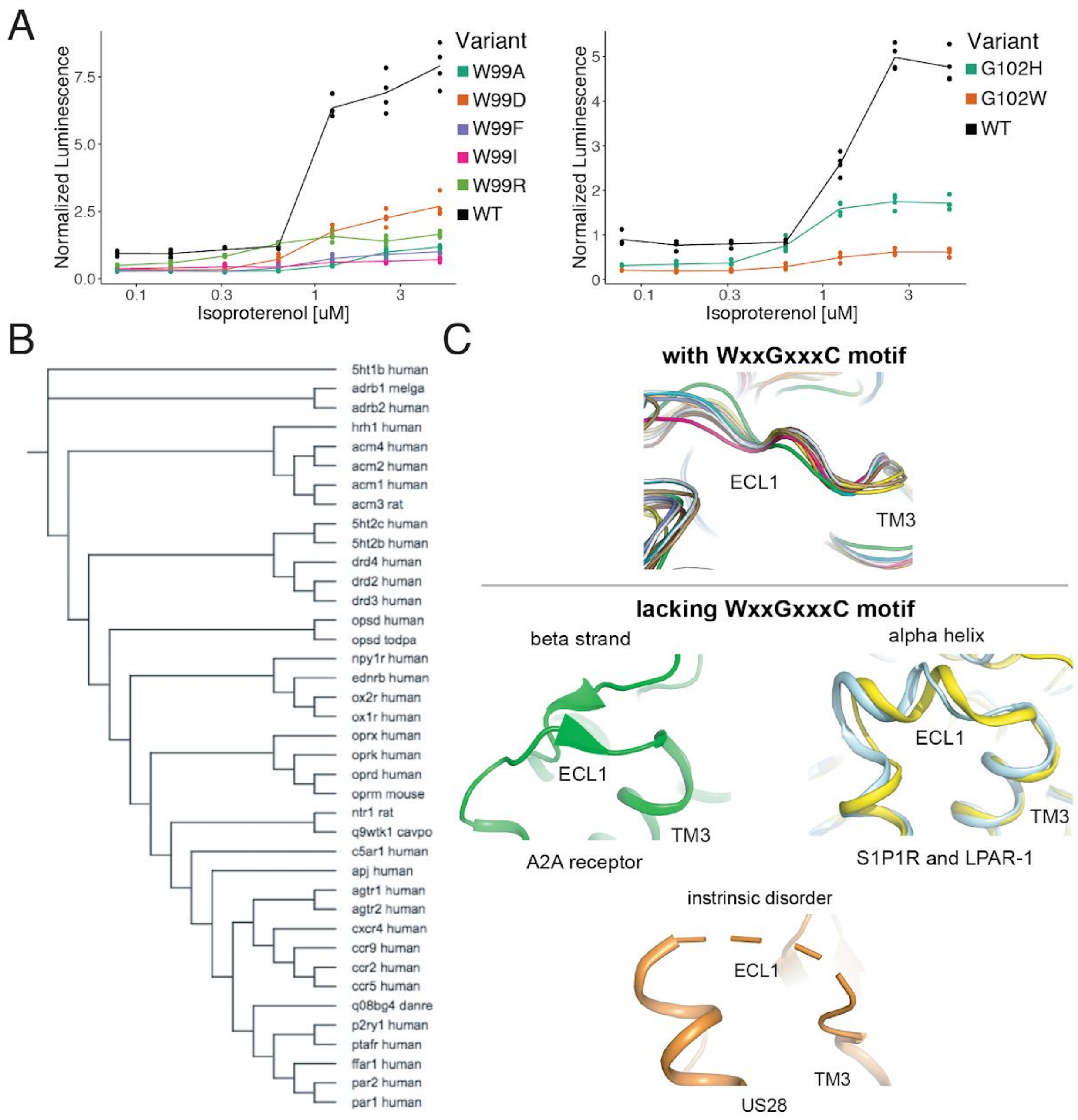


**Supplementary Figure 7. The WxxGxxxC Motif is Highly Conserved Across Class A GPCRs. A.** Individual verification of the mutational intolerance of W99$^{23x50}$ and G102$^{3x21}$. Relative activation of an integrated CRE luciferase reporter gene for β2AR missense variants. **B.** Functional and evolutionary diversity of Class A GPCRs with the WxxGxxxC motif. **C.** ECL1 structures of the four Class A GPCRs lacking the WxxGxxxC Motif, S1P1R, LPAR-1, A2A, and US28.

**Supplementary Table 1. Processed Data.** A table with the processed data used in this study. Includes the position, mutation, min activity, max activity, average activity, propagated uncertainty, coefficient of variation, number of barcodes for that mutation, mutation class, and an annotation of where in the β2AR the position is.

**Supplementary Table 2. Mutational Tolerance.** A table containing positions in the β2AR with their mutational tolerance, rank order, and an annotation for known positions in the receptor.

**Supplementary Table 3. List of Primers used in this Study.** A table documenting the DNA sequences and application of important primers in this study.

### Methods

#### Experimental Methods

##### Endogenous ADRB2 Deletion using CRISPR/Cas9

Cas9 and sgRNAs targeting the sole exon of ADRB2 were cloned and transfected into HEK293T cells according to the protocol outlined in Ran et al. (2013) (Supplementary Table 3)[4]. After transfection, cells were seeded in a 96-well plate at a density of 0.5 cells/well. Wells were examined for single colonies after 3 days and expanded to 24-well plates after 7 days. Clones were screened for ADRB2 deletion by screening them for the inability to endogenously activate a cAMP genetic reporter when stimulated with the ADRB2 agonist isoproterenol. Clones were seeded side by side wild type HEK293T cells at a density of 7,333 cells/well in a poly-D lysine coated 96-well plate. 24 hours later, cells were transfected with 10 ng/well of a plasmid encoding luciferase driven by a cyclic AMP response element and 5 ng/well of a plasmid encoding Renilla luciferase with lipofectamine 2000. 24 hours later, media was removed and cells were stimulated with 25 µl of a range of 0 to 10 µM isoproterenol (Sigma-Aldrich) in CD293 (Thermo Fisher Scientific) for 4 hours.  After agonist stimulation, the Dual-Glo Luciferase Assay kit was administered according to the manufacturer’s instructions. Luminescence was measured using the M1000 plate reader (Tecan). All luminescence values were normalized to Renilla luciferase activity to control for transfection efficiency in a given well. Data were analyzed with Microsoft Excel and R.

##### Landing Pad Genome Editing

The H11 locus was edited using TALEN plasmids received from Addgene (#51554, #51555). HEK293T cells were seeded at a density of 75k cells in a 24-well plate. 24 hours after seeding cells were transfected with 50 ng LT plasmid, 50 ng RT plasmid, and 400 ng of the Linearized Landing Pad using Lipofectamine 2000. 2 days after transfection, cells were expanded to a 6-well plate and one day after expansion 500 ug/ml hygromycin B (Thermo FIsher Scientific) was added to the media. Cells were grown under selection for 10 days. After selection, cells were seeded in a 96-well plate at a density of 0.5 cells/well. Wells were examined for single colonies after 3 days and expanded to 24-well plates after 7 days. gDNA was purified using the Quick-gDNA Miniprep kit (Zymo Research) from the colonies and PCR was performed with Hifi Master Mix to ensure the landing pad was present at the correct locus (LP001F and R). The reaction and cycling conditions are optimized as follows: 95°C for 3 minutes, 35 cycles of 98°C for 20 seconds, 63°C for 15 seconds, and 72°C for 40 seconds, followed by an extension of 72°C for 2 minutes. To ensure a single landing pad was present per cell, HEK293T cell lines with both singly and doubly-integrated landing pads along with untransduced (WT) HEK293T cells were plated at 4 x 10^5 cells per 6-well. All landing pad cells were transfected the next day with 1.094 µg of both an attB-containing eGFP and mCherry donor plasmid and 0.3125 µg of the Bxb1 expression vector or a pUC19 control. Two singly-integrated landing pad cell samples were also transfected with 2.1875 µg of either an attB-containing eGFP and mCherry donor plasmid with 0.3125 µg of the Bxb1 expression vector. Cells were transfected at a 1:1.5 DNA:Lipofectamine ratio with Lipofectamine 3000. 2 days later cells were passaged at 1:10 and were analyzed using flow cytometry 10 days later after 4 total passages. Samples were flown using the LSRII at the UCLA Eli & Edythe Broad Center of Regenerative Medicine & Stem Cell Research Flow Cytometry Core. Cytometer settings were adjusted to the settings: FSC – 183 V, SSC – 227 V, PE-Texas Red – 336 V, Alexa Fluor 488 – 275 V.

#### Individual Donor Bxb1 Recombinase Plasmid Integrations

HEK293T derived cells engineered to contain the Bxb1 Recombinase site at the H11 locus were seeded at a density of 350k cells in a 6-well plate (Corning). 24 hours after seeding cells were transfected with 2 ug Donor plasmid and 500 ng plasmid encoding the Bxb1 recombinase using Lipofectamine 3000 (Thermo Fisher Scientific). 3 days after transfection cells were expanded to a T-75 flask (Corning) and 8 ug/ml blasticidin (Thermo Fisher Scientific) was added one day after expansion. Cells were kept under selection 7-10 days and passaged twice 1:10 to ensure removal of transient plasmid DNA.

#### Ligand-Receptor Activation Luciferase Assay for Genomically Integrated Receptor/Reporter Constructs

HEK293T and HEK293T derived cells integrated with the combined receptor/reporter plasmids were plated at a density of 7333 cells/well in 100 uL DMEM in poly-D-lysine coated 96-well plates. 48 hours later, media was removed and cells were stimulated with 25 µl of a range of isoproterenol concentrations in CD293 for 4 hours. After agonist stimulation, the Dual-Glo Luciferase Assay kit was administered according to the manufacturer’s instructions. Luminescence was measured using the M1000 plate reader. Data were analyzed with Microsoft Excel and R.

#### Ligand-Receptor Activation q-RT PCR Assay for Genomically Integrated Receptor/Reporter Constructs

HEK293T and HEK293T derived cells integrated with the combined receptor/reporter plasmids were plated at a density of 200k cells/well in 2 mL DMEM in 6-well plates. 48 hours after seeding, media was removed and cells were induced with various concentrations of either forskolin (Sigma-Aldrich) or isoproterenol diluted in 1 ml of OptiMEM (Thermo Fisher) per plate for 3 hours. After stimulation, media was removed and 600 uL of RLT buffer (Qiagen) was added to each well to lyse cells. Lysate from each sample were homogenized with the QIAshredder kit (Qiagen) and total RNA was prepared from each sample using the RNeasy Mini Kit with the optional on-column DNAse step (Qiagen). 5 ug of total RNA per sample was reverse transcribed with Superscript III (Thermo-Fisher) using a gene specific primer for the reporter gene and GAPDH (Supplementary Table 3) according the manufacturer's protocol. The reaction conditions are as follows: Annealing: [65°C for 5 min, 0°C for 1 min] Extension: [52°C for 60 min, 70°C for 15 min]. 10% of the RT reaction was amplified in triplicate for both genes, the reporter gene and GAPDH (Supplementary Table 3), using the SYBR FAST qPCR Master mix (Kapa Biosystems) with a CFX Connect Thermocycler (Biorad). The reaction and cycling conditions are optimized as follows: 95°C for 3 minutes, 40 cycles of 95°C for 3 seconds and 60°C for 20 seconds. Reporter gene expression was normalized to GAPDH expression for each sample. Data were analyzed with Microsoft Excel and R.

#### Variant Library Generation and Cloning

The ADRB2 missense variant library was created by splitting the protein coding sequence into 8 distinct segments (~52 a.a. each) and synthesizing all single amino acid substitutions for each segment separately as an oligonucleotide library (Agilent). 500 pg of the oligonucleotide library was amplified with biotinylated primers unique for each segment (Supplementary Table 3) with the Real-Time Library Amplification Kit (Kapa Biosystems) on a CFX Connect Thermocycler (Biorad). The reaction and cycling conditions are as follows: 98°C for 45 seconds, X cycles of 98°C for 15 seconds, 65°C for 30 seconds, and 72°C for 30 seconds, followed by an extension of 72°C for 1 minute. The number of cycles for the amplification was determined to ensure the amplification was in the exponential phase at least two cycles before the amplification reached saturation. The PCR products were cleaned up with the DNA Clean and Concentrator Kit (Zymo Research) and digested with restriction enzymes BamHI and BspQI, BbsI and BspQI, or BbsI and NheI (New England Biolabs). Digestions were cleaned up with the DNA Clean and Concentrator Kit and digested ends of the amplified library were removed by performing a streptavidin bead cleanup with the Dynabeads M-280 and the DynaMag (Thermo Fisher). Each library segment was to be cloned into a different vector that includes components of the ADRB2 reporter and the wild type sequence portion of ADRB2 upstream of the segment being cloned. These eight different base vectors were digested with restriction enzymes BamHI and BspQI, BbsI and BspQI, or BbsI and NheI. The base vectors were cleaned up with the DNA Clean and Concentrator Kit and the library segments were ligated into the base vectors with T4 DNA ligase (New England Biolabs). The ligations were transformed into 5-alpha Electrocompetent cells (New England Biolabs) directly into liquid culture. Cultures were grown at 30° C overnight to maintain library diversity and dilutions were plated on agarose plates to ensure transformation efficiency was high enough to cover the entire library (>100 transformants per library member). DNA was prepared 16 hours later with the DNA Miniprep Kit (Qiagen). The vectors were digested with BspQI and AgeI or NheI and AgeI (Qiagen). Vectors containing unique sequences corresponding to each library segment that complete the ADRB2 protein sequence and reporter were digested with the same restriction enzymes. These fragments were gel isolated from a 1% agarose gel using the Zymoclean Gel DNA Recovery Kit (Zymo Research). These secondary fragments were cloned into the library vectors with the same protocol as the previous cloning step. DNA was prepared 16 hours later with the Plasmid Plus DNA Maxiprep Kit (Qiagen).

#### Variant-Barcode Mapping

After the initial cloning of the variant fragments from the oligonucleotide library into each segment's corresponding base vector, the random barcode attached to each variant was associated to its variant with paired-end sequencing. Each plasmid was amplified with 2 rounds of PCRs with distinct primer sets for each segment (Supplementary Table 3) with HiFi DNA Master Mix (Kapa Biosystems). For the first round of amplification, the reaction and cycling conditions were optimized as follows: 98°C for 30 seconds, 10 cycles of 98°C for 8 seconds, 64°C for 15 seconds, and 72°C for 10 seconds, followed by an extension of 72°C for 2 minutes. These amplicons were gel isolated from a 1% agarose gel using the Zymoclean Gel DNA Recovery Kit. Prior to the second round of amplification, the number of cycles to amplify was determined by performing qPCR with the SYBR FAST QPCR Master Mix (Kapa) on the CFX Connect Thermocycler according to the manufacturer's instructions. The Cq determined from the QPCR plus an addition two cycles was used to as the number of cycles to amplify the libraries for the second round of amplification. For the second round of amplification, the reaction and cycling conditions were optimized as follows: 98°C for 30 seconds, X cycles of 98°C for 8 seconds, 62°C for 15 seconds, and 72°C for 10 seconds, followed by an extension of 72°C for 2 minutes. These amplicons were gel isolated from a 1% agarose gel using the Zymoclean Gel DNA Recovery Kit.  Kit. Library concentrations were quantified using a TapeStation 2200 (Agilent) and a Qubit (Thermo Fisher). The libraries were sequenced with paired end 150-bp reads on a NextSeq 500 in medium-output mode and paired end 250-bp reads on a MiSeq (Illumina).

#### Variant Library Bxb1 Recombinase Plasmid Integrations

HEK293T derived cells engineered to contain the Bxb1 Recombinase site at the H11 locus and deletion of endogenous ADRB2 were seeded at a density of 2.13 million cells per dish in 6 100 mm x 20 mm tissue-culture treated culture dishes (Corning). 24 hours after seeding cells were transfected with 11.5 ug Donor plasmid and 2.9 ug plasmid encoding the Bxb1 recombinase using Lipofectamine 3000. Three days after transfection cells were expanded to T-225 flasks (Corning) and 8 ug/ml blasticidin was added one day after expansion. Cells were kept under selection 7-10 days and passaged 1:10 four times to ensure removal of transient plasmid DNA.

#### Multiplexed Variant Functional Assay Agonist Stimulation, RNA Preparation and Sequencing

HEK293T derived cells engineered to contain the Bxb1 Recombinase site at the H11 locus, deletion of endogenous ADRB2, and integration of the ADRB2 mutagenic library were seeded at a density of 3,237,868 cells per dish in 150 mm x 25 mm tissue-culture treated culture dishes. 10 dishes were seeded for each biological replicate of each drug condition. 48 hours after seeding, media was removed and cells were induced with various concentrations of either forskolin or isoproterenol diluted in 9 ml of OptiMEM per plate for 3 hours. After stimulation, media was removed and 3.24 ml of RLT buffer was added to each well to lyse cells. Lysate from dishes belonging to the same replicate were pooled and vortexed thoroughly. 5 ml of lysate from each sample were homogenized with the QIAshredder kit and total RNA was prepared from each sample using the RNeasy Midi Kit with the optional on-column DNAse step (Qiagen) and eluted into 500 ul $H_2O$. 40 reverse transcriptase reactions were carried out for each sample using the Superscript IV RT kit (Thermo Fisher). For each reaction 11 ul of total RNA were added to 1 ul dNTPs (Qiagen) and 1 ul 2 uM RT primer (Supplementary Table 3). The primers were annealed to the template by heating to 65°C for 5 minutes and cooling down to 0°C for 1 minute. After annealing, 4 ul of RT buffer, 1 ul DTT, 1 ul of RNAseOUT, and 1 ul SSIV were added to the mixture and cDNA synthesis was performed. The reaction and cycling conditions are as follows: 52°C for 1 hour, 80°C for 10 minutes. cDNA from the same sample was pooled together and treated with 100 ug/ml RNAse A (Thermo Fisher) and 200 U of RNase H (Enzymatics) at 37°C for 30 minutes. cDNA was concentrated using the Amicon Ultra 0.5 mL 30k Centrifugal Filter (Millipore) according to the manufacturer's instructions with a final spin step time of 15 minutes. To determine the number of cycles necessary for library amplification in preparation for RNA-seq, 1 ul of cDNA from each sample was amplified with SYBR FAST QPCR Master Mix according to the manufacturer's instructions using primers for library amplification and adaptor addition (Supplementary Table 3). Each sample was subsequently amplified for 4 cycles more than the Cq calculated in the QPCR run adjusting for sample volume. The entire volume of concentrated cDNA for each sample was amplified with sequencing adaptors using NEB-Next High-Fidelity 2x PCR Master Mix (New England Biolabs): 25 ul Master Mix, 2.5 ul of both 10 uM forward and reverse primer (Supplementary Table 3), 4 ul of cDNA, and 16 ul $H_2O$. The reaction and cycling conditions are as follows: 98°C for 30 seconds, X cycles of 98°C for 8 seconds, 66°C for 20 seconds, and 72°C for 10 seconds, followed by an extension of 72°C for 2 minutes. Amplified DNA was purified with the DNA Clean and Concentrator kit and gel isolated from a 1% agarose gel with the Zymoclean Gel DNA Recovery Kit. Library concentrations were quantified using a TapeStation 2200 and a Qubit. The libraries were sequenced with an i7 index read and a single end 75-bp read on a NextSeq 500 in high-output mode.

### Quantification and Statistical Analysis

#### Barcode Mapping

We used the BBTools suite([https://jgi.doe.gov/data-and-tools/bbtools/](https://jgi.doe.gov/data-and-tools/bbtools/)) of programs to process our sequencing data using the default settings unless otherwise noted. First, we used BBDuk2 to filter out any reads matching PhiX (k=23, mink=11, hdist=1) and to trim off any Illumina sequencing adapters. We then used BBMerge to merge our paired end reads. We performed another round of trimming with BBDuk2 to ensure no adapters were left over after merging and to remove any sequence with an N base call. After merging and trimming the reads, we used a custom Python script (bcmap.py) to generate a consensus nucleotide sequence for each barcode.

Briefly the script works as follows. First, we split each read into the 15 nt barcode and its corresponding variant. We then generate a dictionary that maps each barcode to its list of unique sequences and their counts. To enable majority basecalls, we drop any barcode that has less than 3 reads. We then pass the barcodes through a series of filters to eliminate potential errors introduced by barcodes that are mapped to multiple variants. Since we barcoded and mutagenized the ADRB2 gene in separate pieces, barcodes can be contaminated with variants from different parts of the ADRB2 gene.

We address this case by using BBMap to align every barcode's sequences to the ADRB2 reference and consider that barcode to be contaminated if any sequence aligns >5 nt away from the most common sequence. Another source of contamination comes from the chip-synthesized library itself, which contains a significant number of single base deletions. We consider a barcode contaminated if it has any sequences of different lengths as it is unlikely that a single base deletion will come from an Illumina sequencer by chance. However, these filters would not catch the case where a barcode is contaminated with variants from the same piece of ADRB2. As we only synthesized the missense variants, we expect variants within the same piece of ADRB2 to be a Levenshtein distance of 4 from each other on average (approximately two changes to WT and two changes to a new codon). Thus, we drop any barcode that has a sequence with >1 read at a Levenshtein distance of 4 away from that barcode's most common sequence. Lastly, we generate a consensus sequence by taking the majority base call at each position and call an N at any ties.

After we associate each barcode with its consensus sequence, we use a series of different alignments to determine that sequence's identity. To find the designed missense variants in our library, we use BBMap to search for barcodes that have an exact alignment to them. To find frameshift mutations, we use BBMap to align the consensus sequences to the ADRB2 reference and parse the resulting CIGAR strings for indels with a simple python script (classify-negs.py). Finding synonymous mutants required more processing as each sub-library did not start at a complete codon. We first used the rough BBMap alignment to determine what ADRB2 chunk each sequence was associated with. We then used a custom python script (synon-filter.py) to trim up to the last whole clonal codon, as the first few codons of each sequence were part of the clonal backbone and are unlikely to have any errors. Finally, we translated the resulting sequences, aligned the protein sequence to the ADRB2 coding sequence with a Smith-Waterman aligner from the Parasail library (https://github.com/jeffdaily/parasail), and retained perfect translations with the correct length.

#### Data Normalization

We incubated our cellular library with forskolin to activate the cAMP reporter in each cell, providing an agonist-independent measurement of maximal reporter activity. This measurement can be used to approximate cellular copy number. To ensure that barcodes with low cellular representation are excluded from our analyses, we require all barcodes to be present in both forskolin repeats, and filter out any barcodes with a mean reads per million less than 0.2 (~8-10 reads at our sequencing depth). We also excluded barcodes with high forskolin counts (>= 10 RPM) as they are systematically less induced in the drug conditions relative to other barcodes. Next, we require that all of the barcodes in the forskolin condition are also present in our drug conditions, and set any missing barcodes to 0. We then add a pseudocount that is scaled relative to the condition with the fewest number of reads (N/min(N)), and normalize each condition to its read depth (including added pseudocounts)[5]. Finally, we divide this value by its associated forskolin value to control for variation in cellular abundance.

Since each variant in our library was associated with a median of 10 barcodes, we took the average of all barcodes. We then defined activity as the ratio of these values to the value of the mean frameshift. Finally, we averaged the relative activities of our two repeats together and used propagation of uncertainty to combine their standard deviations.

#### Conservation, EVMutation, and gnomAD

To calculate sequence conservation at a species level, we aligned 55 ADRB2 orthologs from the OMA database (entry: HUMAN24043) using MAFFT with the default settings (mafft --reorder --auto). For Class-A GPCRs, we retrieved the multiple sequence alignment from GPCRdb. We then used the Jensen-Shannon Divergence[6–8] to score both of these alignments. We only considered conservation scores at positions in the MSA that contained residues from the β2AR. For both EVMutation and gnomAD, we simply downloaded the results for ADRB2. We considered residues in gnomAD to be potentially loss of function if their mean activity plus the standard error of the mean was less than two standard deviations from the mean of the frameshift distribution.

#### Unsupervised Learning

We performed a number of preprocessing steps before running UMAP on our data. First, we grouped amino acids into 8 different classes based on their physicochemical properties ((+) - R, H, K; (-) - D, E; Aromatic - F, W, Y; Amide - N, Q; Nucleophilic - C, S, T; Hydrophobic - I, L, V, M; Small - G, A; Proline - P) and averaged their relative activities. Next, we standardized the log2 relative activity values of each group and used mean imputation to model missing data for any missing AA groups at a given position. Finally, we combined the data from every drug condition into a 412 x 32 design matrix in which the columns are an AA group at a specific condition and the rows are the positions in the protein.

With our data processed, we used the R implementation of UMAP to run hyperparameter search (https://github.com/jlmelville/uwot) of all combinations of UMAP embeddings with the parameters n_neighbors = (4, 8, 16, 32) and n_components = (2,

3, 4, 5, 6, 7, 8, 9, 10), holding min_dist=0 and n_epochs=2000 constant. This provided a variety of different representations of our data that we used HDBSCAN[2] to search for clusters in these embeddings (R package dbscan; minPts = 10). To ease interpretation of the clustering, we plotted the HDBSCAN results onto a 2D UMAP embedding with the following parameters: n_neighbors=4, min_dist=0, n_components=2, n_epochs=2000, and random_state=3308004 using the Python implementation[1] (https://github.com/lmcinnes/umap). We found the cluster assignments to be largely robust across the different embeddings, and used them to guide our manual cluster assignment.

#### Structural Modeling and Solvent Accessible Surface Area

Molecular graphics and analyses were performed with UCSF Chimera[9] and PyMol. To determine if a given position in the $\beta_2$AR points into the core of the protein or into the lipid membrane, we used FreeSASA[10] (version 2.0.3) to calculate the Solvent Accessible Surface Area (SASA) of the $G_s$-bound $\beta_2$AR (PDB: 3SN6). The $G_s$ occludes the intracellular surface of the $\beta_2$AR thereby reducing the SASA of residues on the intracellular surface. Similarly, the extracellular surface is mostly blocked by the extracellular loops. Finally, we used the Orientations of Proteins in Membranes (OPM) database[11] to filter out any residues outside of the lipid membrane from our analyses. To quantify charge sensitivity, we calculated the average activity for H, K, R, D, and E substitutions at each agonist concentration for residues in the lipid membrane. We then multiplied the values by -1 and standardized the results within each agonist concentration group such that the values were mean-centered and scaled by their standard deviation. We calculated hydrophobic sensitivity (I, L, V, M) in an analogous manner. Next, we classified residues that had above average charge sensitivity and below average hydrophobic sensitivity as being exclusively charge sensitive. Conversely, we classified residues that had above average charge sensitivity and above average hydrophobic sensitivity as being intolerant.

#### Structural and Sequence Analysis of Class A GPCRs

For the structural analysis, the crystal structures of class A GPCRs were obtained from PDB[12]. In order to compare structures across the different sub-families of class A GPCRs, structures of representative examples from each subfamily were chosen. In order to compare structures across the conformational states in different GPCRs, the structures of beta-2 adrenergic receptor, M2 muscarinic receptor, kappa-opioid receptor, and mu-opioid receptor were chosen. A2A receptor and mammalian rhodopsin were excluded as they lacked a Trp residue in the first extracellular loop. These receptors were chosen due to the availability of pairs of inactive state and active state structures. The inactive state structures were identified based on the presence of co-crystallized antagonist/inverse-agonist and the active state structures were identified based on the presence of a co-crystallized agonist and co-complexed interacting partner at the G protein-coupling site. Structural alignment and measurement of inter-atomic distances were performed using PyMOL (https://pymol.org). The structure alignment was performed over the sequence stretch between the conserved Trp/Phe residue in the first extracellular loop and the canonical disulfide bridge forming Cys3x25 (GPCRdb number) present on TM3.

For the sequence analysis, the sequence alignment of class A GPCRs was obtained from GPCRdb[13]. The alignment was filtered for receptors that contained the canonical disulfide bridge forming on TM3 residue in ECL1, which gave a total of 202 GPCRs sequences. Using this alignment, the sequence logo was made using the Weblogo program (https://weblogo.berkeley.edu/logo.cgi).

#### Statistical Tests

All statistical tests unless otherwise noted are the two-sided Mann-Whitney U test and were performed in R (version 3.5.x) using the wilcox.test function.

#### Software

All code is available at https://www.github.com/KosuriLab/b2-dms. Sequencing data can be accessed from the sequencing read archive (SRA) with the accession number XXXXXXXXX. To avoid potential visual distortions in the heatmap, we used perceptually uniform color maps[14]. For parallelization, we employed GNU Parallel[15].
